# Hierarchical processing and polarization encoding in the cephalopod visual system

**DOI:** 10.64898/2026.01.08.698312

**Authors:** Tomoyuki Mano, Konstantinos Tsaridis, Yutaka Kojima, Giovanni D. Masucci, Thi Thu Van Dinh, Rudi Tong, Vasileios Glykos, Lada Dolezalova, Keishu Asada, Daria Shumkova, Loranzie S. Rogers, Maxime Hamon, Matteo Santon, Makoto Hiroi, Teresa L. Iglesias, Nicholas W. Bellono, Martin J. How, Yukiko Goda, Leenoy Meshulam, Sam Reiter

## Abstract

Coleoid cephalopods (octopus, cuttlefish and squid, hereafter ‘cephalopods’) have evolved a range of complex visually-guided behaviours, from dexterous hunting to skin-pattern based camouflage and communication^1^. They have also evolved sensitivity to the polarization of light^2^, an adaptation thought to help detect camouflaged or semitransparent predators and prey in low visibility underwater environments^3–10^. How visual information is processed by the cephalopod brain to support their behaviours remains unclear. Here, studying the bigfin reef squid *Sepioteuthis lessoniana*, we performed calcium imaging and electrophysiological recordings from populations of neurons in the large visual center of the cephalopod brain, the optic lobe (OL). We revealed that the retina-recipient superficial OL contains a diversity of functionally distinct cell types, spatially organized into sub-layers, processing spatio-temporal features of light intensity and possessing polarization angle specificity. More complex features, e.g. direction selectivity, are seen in deeper regions of the OL cortex, which also exhibits spontaneous waves of neural activity in the absence of visual input. Neurons in the downstream OL medulla exhibit visual receptive field sizes and spontaneous activity levels which increase with brain depth, consistent with the hierarchical processing of visual information through the medulla’s tree-like anatomical organization. Medulla neurons exhibit sensitivity to local decreases in the degree of linear polarization (DoLP), which they integrate additively with light intensity information. Underwater imaging in the squid’s habitat off the coast of Okinawa, Japan, demonstrate that polarization sensitivity confers a robust short-range boost in object-background contrast over a range of objects and environmental conditions. These findings reveal convergent principles of hierarchical visual processing shared between cephalopods and vertebrates, and highlight how cephalopods utilise their distinct adaptation of polarization sensitivity to solve universal visual challenges underwater.

## Main

Like the vertebrate eye, the cephalopod eye focuses light through a lens onto a retina containing photoreceptor cells^11^ (Fig. 1a,b). Unlike in vertebrates, cephalopod photoreceptors are intrinsically sensitive to the angle of linear polarization of light^12^, with microvilli spatially arranged so as to respond maximally to either vertically or horizontally polarized light stimuli (Fig. 1c). Photoreceptors send axons directly in small bundles to the central brain, where they synapse onto the outer region of the ipsilateral optic lobe (OL cortex)^13,14^. Cajal termed this area the ‘deep retina’, suggesting a functional analogy based on anatomical similarities to the vertebrate retina (Fig. 1d)^13–15^. The OL cortex is organised retinotopically^16^, and ex vivo recordings have demonstrated layer-dependent hotspots sensitive to different spatiotemporal features of light onset or offset, indicative of OL visual computation^17^. The underlying OL medulla is composed of ‘islands’ of densely packed neurons separated by neuropil, with island size increasing with distance from the brain surface^18^. Behavioural studies have demonstrated that cephalopods perceive changes in the polarization of light^19,20^, a sensitivity thought to aid detection of predators and prey, particularly in underwater environments subject to caustic-intensity flicker^3,4^ and turbid, low-visibility conditions^5–7^ that erode intensity contrast even at short viewing distances. Notably, cuttlefish and squid also produce skin patterns which reflect polarized light, and may be used as a communication channel invisible to most observers^21–26^. Neural computations in the cephalopod brain remain unclear, in part due to difficulties in making neural recordings from live animals. Here, by developing methods for *in vivo* neurophysiology in cephalopods at cellular resolution, we investigated directly how neural populations at different levels of the OL transform light intensity and polarization information into formats that support visual behaviour.

**Figure 1.**
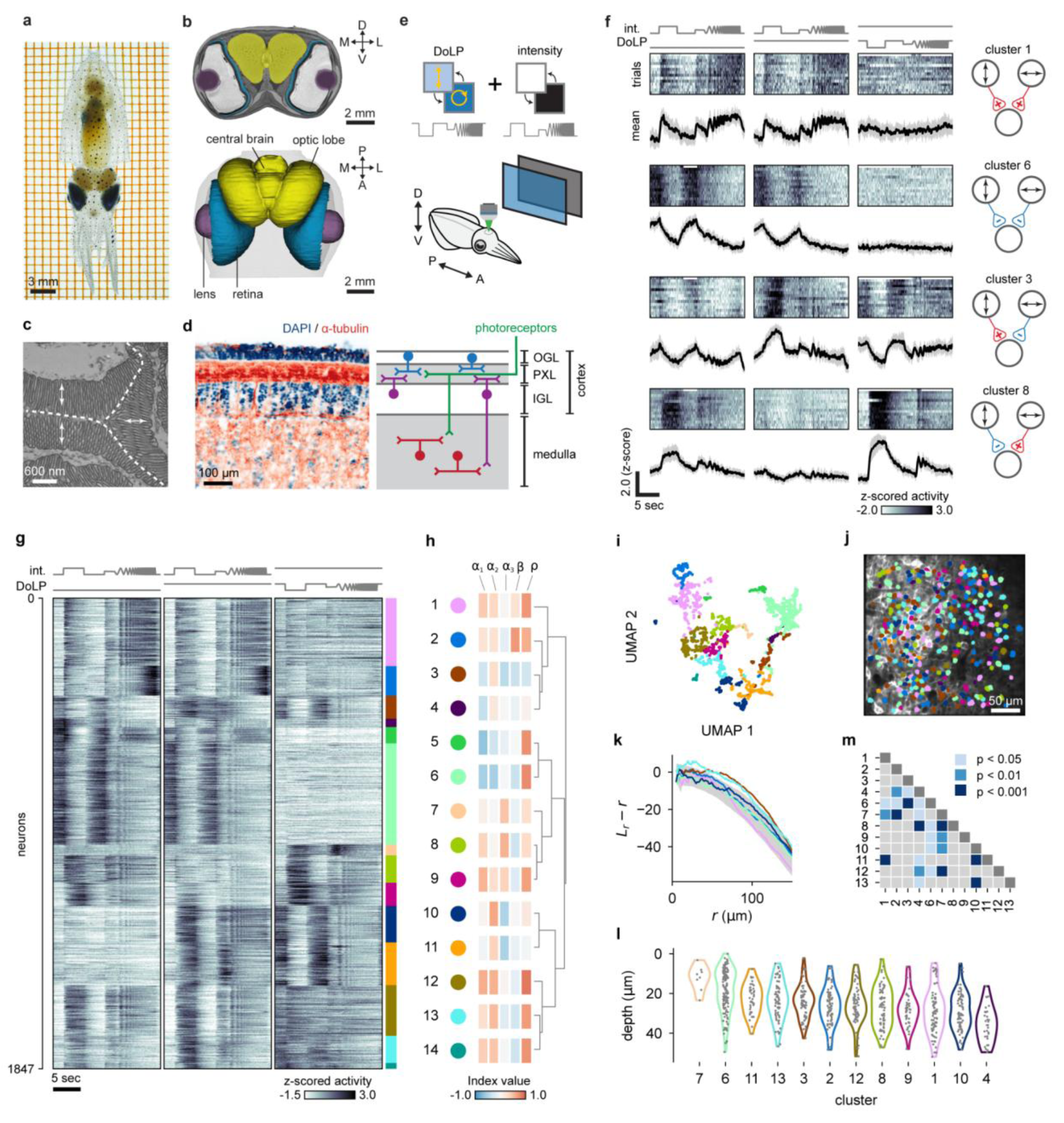
| Functionally distinct cell types of the OL cortex. **a**, Juvenile *Sepioteuthis lessoniana*. **b**, MRI scan of the squid head, with segmented brain and eyes (Methods). **c**, Transmission electron microscopy image through squid photoreceptors. Arrows show orthogonal arrangement of microvilli in different cells, supporting polarization angle sensitivity. **d**, Slice through the superficial OL (DAPI: nuclear stain, α-tubulin: neuropil marker). Right: schematic of brain regions and general neuroanatomy. **e**, Schematic of the full-field visual stimulation experiment with controlled light intensity and DoLP. **f**, Example Ca^2+^ responses from four cells. Right: synaptic input from photoreceptors, inferred from patterns of visual responses. Clusters as in h. **g**, The landscape of neural Ca^2+^ responses to intensity and DoLP changes, pooled dataset of OGL (n = 1085 neurons, 5 animals) and IGL (n = 762 neurons, 8 animals). Colours: cluster identity. **h**, Cluster-wise “barcode” response summary. α₁, α₂, α₃: activation indices for phases 1-3; β: high-frequency modulation index; ρ: correlation between responses in phases 1 and 2. Dendrogram: similarity relationships between clusters (Methods). **i**, UMAP embedding of all neurons, coloured by cluster identity. **j**, Spatial distribution of functional cell types in a representative OGL recording, with neurons coloured by cluster ID (n = 306 cells). **k**, Ripley’s L function, *L_r_*, for randomness of spatial distribution for neuron clusters 1, 2, 3, 6, 10, and 13 present in j (Methods). *L_r_* – *r* is plotted against *r*, where *r* is a radial distance. Gray shading indicates the range expected from complete spatial randomness (CSR). **l**, Cluster-wise distribution of neurons over OGL depth (0 = brain surface; n = 792 cells, 4 animals). **m**, Pairwise statistical significance map of depth localisation (Dunn’s test with Holm–Bonferroni correction for multiple comparisons).

### Functionally distinct cell types of the OL cortex

We developed techniques for cellular-resolution two-photon calcium imaging from awake, head-fixed squid (*Sepioteuthis lessoniana*), using calcium indicators^17^ (Methods, Extended Data Fig. 1, Supplementary Video 1). We used this method to record from neural populations within the outer and inner granular layer of the OL cortex (OGL, IGL). In vitro patch-clamp electrophysiology combined with cable theory modelling indicated that somatic signals faithfully report these neurons’ overall activity (Methods, Extended Data Fig. 2). To study neural processing of light intensity and polarization, we back-projected images from a projector onto a modified liquid crystal display (LCD) placed parallel to the ipsilateral eye, allowing for the independent variation of light intensity and DoLP patterns (Fig. 1e, Extended Data Fig. 3, Methods)^27^. Eye movements were tracked using high-speed cameras and IR lighting. Squids displayed minimal eye movements under these conditions (Extended Data Fig. 1f). We utilised a full-field visual stimulus consisting of periods of on, off, and a temporal chirp. This stimulus was presented in three interleaved conditions: 1) as changes in light intensity with constant minimal DoLP (0.003), 2) as changes in light intensity with constant maximal DoLP (0.95), and 3) as changes in DoLP (ΔDoLP) with a constant maximal intensity. We presented 10 to 15 trials of each condition, and calcium activity traces from cell bodies were trial averaged to determine cellular responses.

We observed a functional diversity of neurons within the OL cortex. Visual responses varied in both polarity and magnitude to changes in light intensity, ΔDoLP, and temporal features of each. For example, we identified a simple ON cell type, activated by increases in light intensity regardless of DoLP level, and unresponsive to ΔDoLP (cluster 1, Fig. 1f). Similarly, we found polarization-insensitive ON-suppression cells (cluster 6). In another kind of neuron, responses were suppressed by increases of unpolarized intensity, activated by increases in polarized intensity, and suppressed by decreases in DoLP (cluster 3). There were also neurons that behaved in the opposite manner (cluster 8). These response types suggest that the two channels of retinal polarization sensitivity interact within the brain to produce both polarization-insensitive and polarization-opponent neurons.

To classify cell types based on their functional signatures, we clustered neural activity across stimulus conditions using Leiden community detection (Fig. 1g-i, Extended Data Fig. 4, n = 1847 neurons; Methods). UMAP embedding suggested that the tuning features of the neurons were distributed along a continuum, giving rise to different tuning patterns as one traverses along a path in high-dimensional feature space (Fig. 1i). By manually curating algorithmically derived clusters, we identified 14 putative cell types. We further distilled prominent differences across cell types into five parameters (Figure 1h; Methods). Clusters could be differentiated by their activation/suppression in response to light increases or DoLP decreases (α_1_, α_2_, α_3_), activation/suppression to high-frequency stimuli (β), and the correlation of their responses to unpolarized vs. polarized light (ρ).

The spatial mapping of cell types revealed a finely interleaved pattern over the OGL surface (Fig. 1j). To quantify this, we computed Ripley’s L function for each cell type (Methods) within the same imaging plane and found that the traces were consistent with a spatially random distribution (Fig. 1k). A k-th nearest neighbour entropy measure produced similar results (Methods). However, the distribution of functionally defined cell types along the OGL depth was not homogeneous (H = 129.6, p = 2.06 × 10⁻²², Kruskal–Wallis H-test), and a subset of cell types occupied distinct depths within the OGL (Fig. 1l,m, Methods). This functional gradient within the OGL is consistent with reports of sub-laminar structure identified morphologically and transcriptionally^13,28^.

### Comparing functional properties across the OL cortex

OGL and IGL neurons produced similar responses to our full-field temporal stimulus (Extended Data Fig. 4d,e). We therefore delivered a richer stimulus set to investigate the differences between layers. We first used sparse noise to map receptive field position and size. Receptive fields were arranged retinotopically parallel to the OL surface (Extended Data Fig. 5). Receptive field sizes were significantly larger in the IGL than in the OGL (OGL: 33.6 ± 11.8 visual degree^2^, n = 394 neurons, 3 animals; IGL: 44.6 ± 14.4 visual degree^2^; n = 297 neurons, 2 animals; p = 1.03 × 10^-24^, Welch’s t-test), suggesting that IGL neurons integrate multiple inputs before projecting downstream.

To test polarization angle specificity, we delivered a full field flash stimulus while placing a linear polarizer at multiple angles (0-180°, Methods). Visually responsive cortical neurons were either sensitive to horizontal angles, sensitive to vertical angles, or responded to all polarization angles (Extended Data Fig. 6a-g). As expected from a two channel polarization system^29^, polarized intensity stimulation at 45° offset from peak sensitivity evoked responses similar to unpolarized stimuli (Extended Data Fig. 6h,i). DoLP stimulation at this angle evoked weak responses (Extended Data Fig. 6j).

These results motivated a closer examination of responses to vertically vs horizontally polarized light. We delivered our full-field stimulus twice, with a second session following a 90° rotation of the LCD. Some neurons showed opposite response patterns across sessions, indicative of polarization angle-specificity (e.g., Fig. 2a, neuron #1). Other neurons, while sensitive to DoLP stimuli, responded similarly across sessions (e.g., Fig. 2a, neuron #2). Clustering analysis, pooling OGL and IGL neurons, identified 12 putative functional clusters (Fig. 2b, Extended Data Fig. 7a-c; OGL: n = 214 neurons, 4 animals; IGL: n = 297 neurons, 3 animals). Many clusters showed anti-correlated responses to DoLP changes across sessions, quantified by a low degree index (DI). Most responses to DoLP stimuli in both OGL and IGL are therefore specific to polarization angle (Fig. 2c). However, some clusters (e.g., 3, 9, and 10) exhibited polarization angle-invariant responses, characterized by a positive DI. These clusters were predominantly composed of IGL neurons. Indeed, the IGL contained more neurons with strongly positive DI values (DI > 0.3) than the OGL (p = 0.019, one-sided Fisher’s exact test), suggesting that angle-invariant visual responses begin to emerge within the IGL.

**Figure 2.**
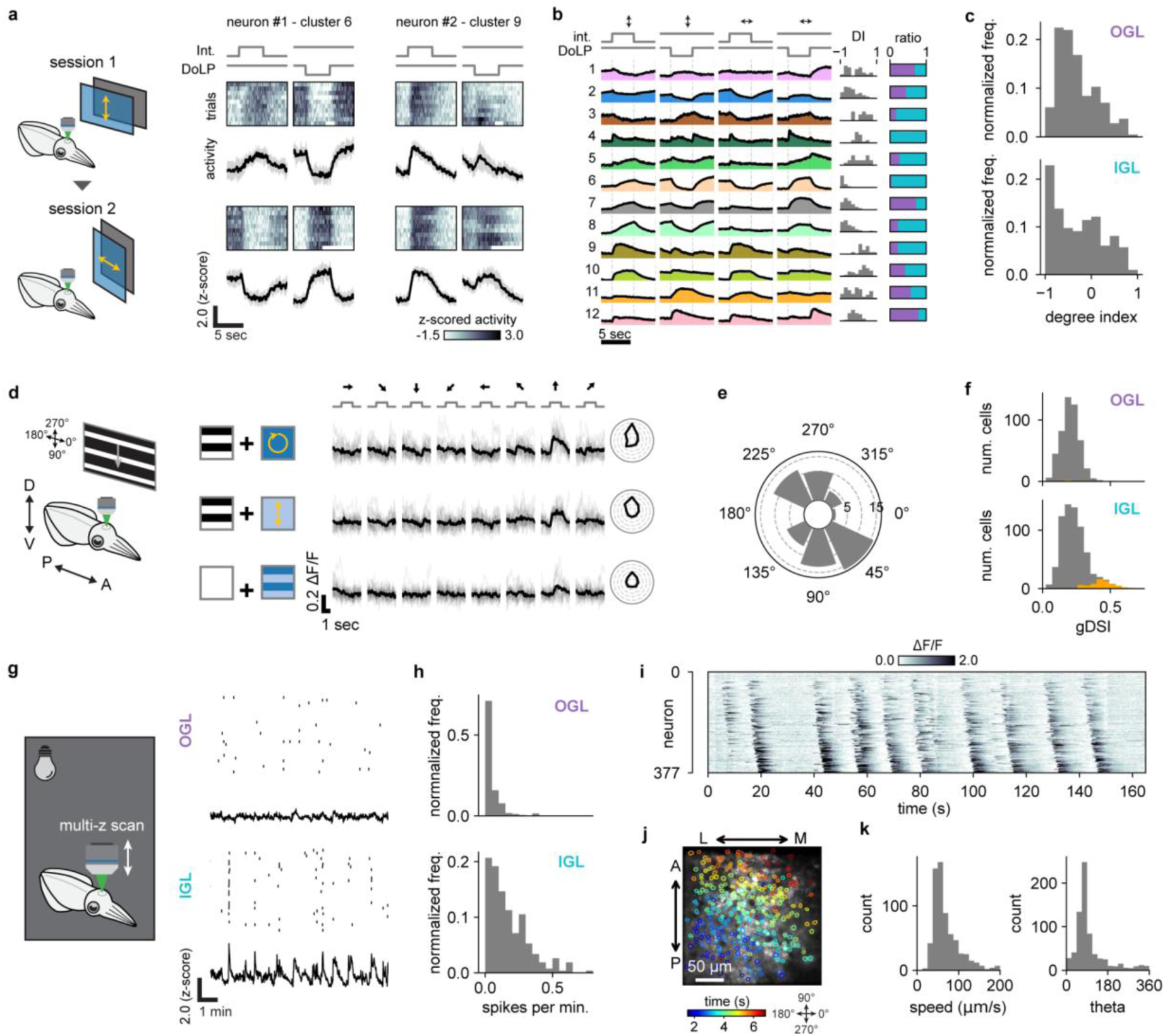
| Comparing functional properties across the OL cortex. **a,** Left: Monitor flip experiment. Right: Example neural Ca^2+^ responses for vertically polarized screen (top rows) and horizontally polarized screen (bottom rows). **b,** Left: Pooled OGL and IGL cluster-averaged Ca^2+^ responses to vertical (first two columns) and horizontal (last two columns) screen orientations. Middle: Degree index (DI) distributions. Right: Ratio of OGL (purple) and IGL (blue) neurons per cluster. **c,** Degree index values (OGL: n = 214 neurons, 4 animals; IGL: n = 297 neurons, 3 animals). **d**, Left: Drifting grating experiment. Right: An example direction-selective (DS) neuron. Top row: intensity grating, unpolarized background. Middle row: intensity grating, vertically polarized background. Bottom row: DoLP grating, constant-intensity background. **e**, Distribution of preferred directions among DS cells in the IGL (n = 63 neurons). **f**, Direction selectivity indices (gDSI) for neurons in the OGL (top) and IGL (bottom). Orange bars: statistically significant direction tuning (OGL: n = 4/491 neurons, 3 animals; IGL: n = 63/695 neurons, 4 animals). **g,** Left: Fast multi-z plane imaging was performed without visual stimuli. Right: Raster plot showing calcium spikes, simultaneously recorded from OGL (top) and IGL (bottom) (n = 30 randomly selected neurons). Bottom trace: average Ca^2+^ signal across all neurons in each imaging plane (OGL: n = 103 neurons, IGL: n = 100 neurons). **h,** Spontaneous calcium activity rates (n = 335 OGL and n = 266 IGL neurons, 3 animals). **i,** Ca^2+^ signals from IGL neurons during a wave bout, sorted by wave onset time. **j,** Propagation of the second wave in i (at approx. 20 s). Cells are colour-coded by peak time. **k,** Distributions of wave propagation speed and direction (pooled n = 3 recordings).

We next investigated the selectivity of neurons to visual movement, a fundamental operation across early visual systems^30^. We delivered drifting grating stimuli moving in eight different directions, with grating contrasts yielded by three intensity/DoLP conditions (Fig. 2d, Methods). We observed a subset of neurons in the IGL sensitive to movement orientation (15.7%) or direction (9.1%) (Fig. 2d-f; Extended Data Fig. 7d-f), using common indices for selectivity (Methods; n = 695 cells, 3 animals). In contrast, orientation or direction selectivity was very rare in the OGL, and occurred significantly less than in the IGL (2.8% and 0.8%, respectively; n = 491 cells, 4 animals). Among direction-selective cells in IGL, 9.5% responded to motion of both light intensity and polarization, with similar directional tuning. Across the IGL population, tuned directions were biased to vertical over horizontal (Fig. 2e).

Simultaneous OGL/IGL imaging via fast z-scanning (Methods) revealed that spontaneous neural activity also differed greatly between the OGL and IGL. In complete darkness, OGL neurons were largely silent (0.04 ± 0.06 spikes per minute, per neuron). In contrast, under the same conditions IGL neurons were often highly active, frequently in synchrony (Fig. 2g,h, 0.17 ± 0.14 spikes per minute, per neuron; p = 1.02 × 10^-32^, Welch’s t-test). Strikingly, we observed waves of neural activity traveling across the OL cortex, restricted to the IGL (multi-plane imaging, 0 waves seen in OGL during 44 IGL waves, 5 animals). IGL waves often occurred as singletons (n = 105 wave events, ∼16 hours of recording, 13 animals). However, in rare instances waves occurred repeatedly in bouts of approximately three minutes (Fig. 2i,j; 3 bouts detected from ∼16 hours of recording). During these bouts, waves propagated predominantly from a posterior to anterior direction, corresponding to downward motion in visual space, at a speed of 72.9 ± 40.6 µm/sec (Fig. 2k). Behaviourally, wave bouts were accompanied by increased breathing rate and erratic twitching behaviours (Supplementary Video 2).

### A visual hierarchy within the OL medulla

We next sought to understand visual processing downstream of the OL cortex, in the medulla and peduncle lobe (PDL). To study the medulla’s peculiar neuroanatomy, we used light sheet imaging to visualise the squid brain in 3D, and constructed a digital atlas through manual segmentation. As in cuttlefish^18^ and octopuses^31^, we observed that the ‘islands’ of densely packed cell bodies visible in brain slices connect to each other, forming a three-dimensional structure resembling a tree (Fig. 3a, Supplementary Video 3). Computational registration of multiple brains revealed that the precise arrangement of tree ‘branches’ differed across individuals (Fig. 3b., 31.5 ± 0.5% IoU similarity score, n = 6 comparisons from 4 brains, Methods). We estimated the local volume of individual 3D tree branches across the full OL using morphological operations (Methods). The average branch volume for small voxels grew as a function of distance from the OL cortex (OL depth) through most of the medulla, before decreasing in size as the neuropil split into discrete bundles upon approaching the central brain. Across individuals, branch volume increased with distance at similar rates (Fig. 3c), pointing to a conserved developmental rule underlying the medulla’s tree-like architecture.

**Figure 3.**
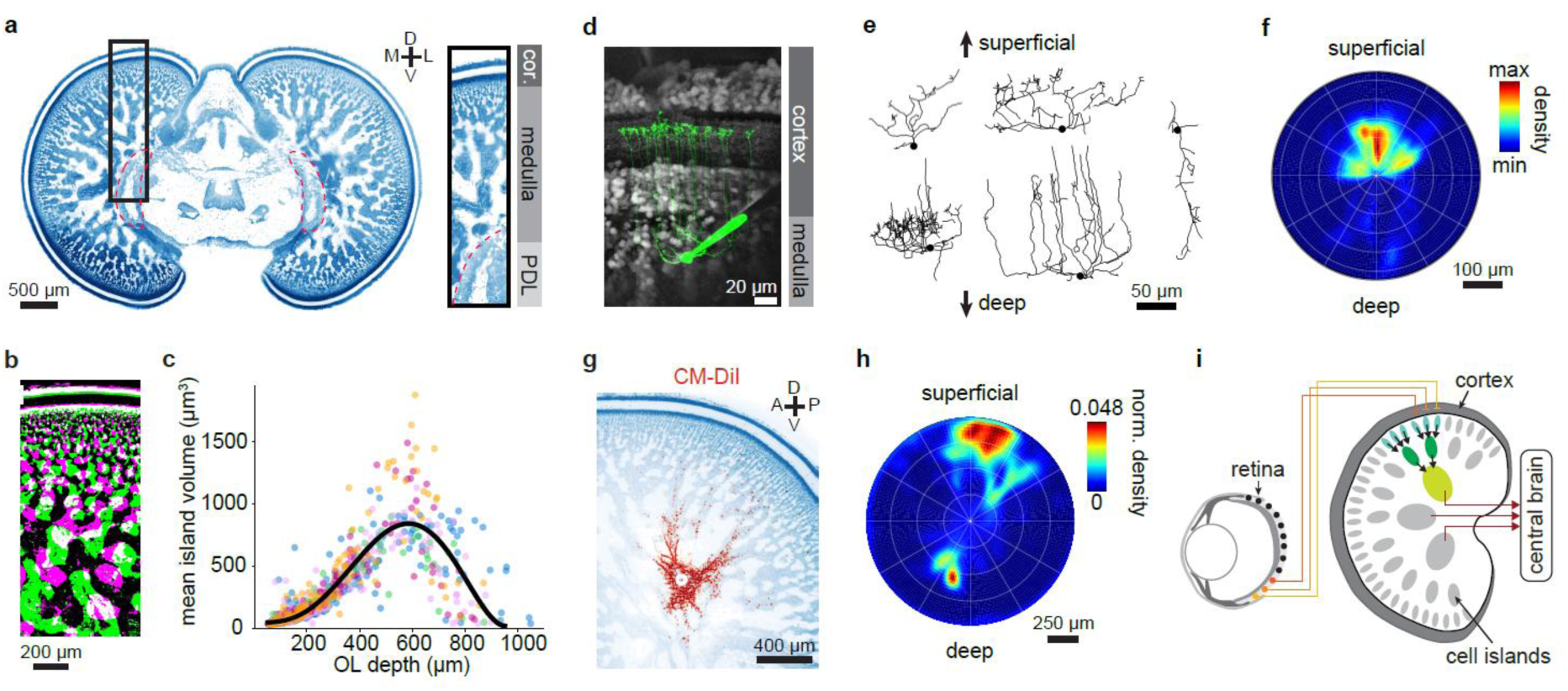
| Hierarchical organization of medulla anatomy. **a,** Squid coronal brain section digitally resliced from 3D light-sheet imaging (nuclear stain) Inset: zoomed view on the OL. Red dashed outline: PDL. **b**, Digital slice of OLs from two animals, following 3D registration. Magenta: animal 1, Green: animal 2. White: overlap. **c**, Growth in local cell island volume with depth in the medulla following 3D registration. Colours: voxels from different brains (n = 5 brains). **d**, Pipette containing 0.1 mM Alexa Fluor 488 filling a medulla neuron, during whole-cell patch clamp. **e**, Example filled and reconstructed neurons, showing processes which extend along the superficial-deep OL axis. **f**, Histogram of neural processes, centered at soma location and rotated to align the superficial-deep OL axis in different slices (n = 19 neurons). **g**, Digital slice of the OL following CM-DiI injection. Circle: injection site. **h**, Radially normalized density of DiI-labelled somata from brain in g. The injection site is at the center, and the vertical axis represents the superficial-to-deep OL axis (Methods). **i**, Schematic of proposed hierarchical flow of visual information through the OL.

To resolve the morphology of individual neurons within the medulla, we cut slices of the OL and made whole-cell patch-clamp recordings to fill neurons with dye (Fig. 3d, Methods). Neural processes in the medulla projected towards or away from the OL surface. From the soma, neurons extended processes perpendicular to the major axis of the local cell island to enter the neuropil. Processes branched towards or away from the OL surface^13^, but rarely in both directions at the same time, tightly following the medulla tree structure (average length = 146.7 ± 16.7 µm, n = 19, Fig. 3e). Neurons with processes extending towards the surface tended to span, and presumably integrate activity from, four to six cell islands (average width = 178.2 ± 15.1 µm, average length = 141.7 ± 14.5 µm, n = 9, Extended Data Fig. 8a). Neurons projecting towards the deep medulla stayed close to the cell island which they originated from but had longer processes (average width = 136.29 ± 44.01μm, average length = 261.25 ± 18.75 µm, n = 3, Extended Data Fig. 8b). Lastly, some neurons stayed contained within a single cell island, likely functioning as local processing units (Extended Data Fig. 8c). Overall, superficially extending processes were much more extensive than those projecting deeper (Fig. 3f).

To study macroscopic connectivity within the medulla tree, we used the tracer CM-DiI to label neurons passing through a local injection site, predominantly via retrograde transport (Methods). 3D light-sheet imaging revealed labelled cell bodies located superficial to the injection site, consistent with neural processes moving along the neuropil separating tree branches (Fig. 3g,h). Outside of the ipsilateral OL, DiI-labelled somata were found in the contralateral OL, and a collection of central brain structures^16^ (Extended Data Fig. 9). The medulla anatomy thus suggests a hierarchy of visual information processing, with neurons receiving polysynaptic input from increasingly large areas of the OL cortex as one moves deeper into the structure (Fig. 3i).

To test this possibility, we adapted our preparation to record from populations of single units throughout the depth of the OL using Neuropixels probes (Extended Data Fig. 10), and employed a registration pipeline to localize recordings in three dimensions (Fig. 4a,b, Extended Data Fig. 11). We used sparse noise visual stimuli to map neural receptive fields (single units, Methods). Medulla neurons were mostly activated by light onset, and often possessed weaker, larger receptive fields to light offset (98% ON neurons, 2% OFF neurons, total: n = 355 neurons with receptive fields on screen, 36 animals). Many neurons were sensitive to moving visual stimuli, exhibiting similar biases to vertical movement as observed in the IGL (Extended Data Fig. 12). Retinotopy was maintained across medulla neurons, demonstrated by a systematic relationship between electrode location and receptive field position in single recordings (Fig. 4c) and overall (Fig. 4d).

**Figure 4.**
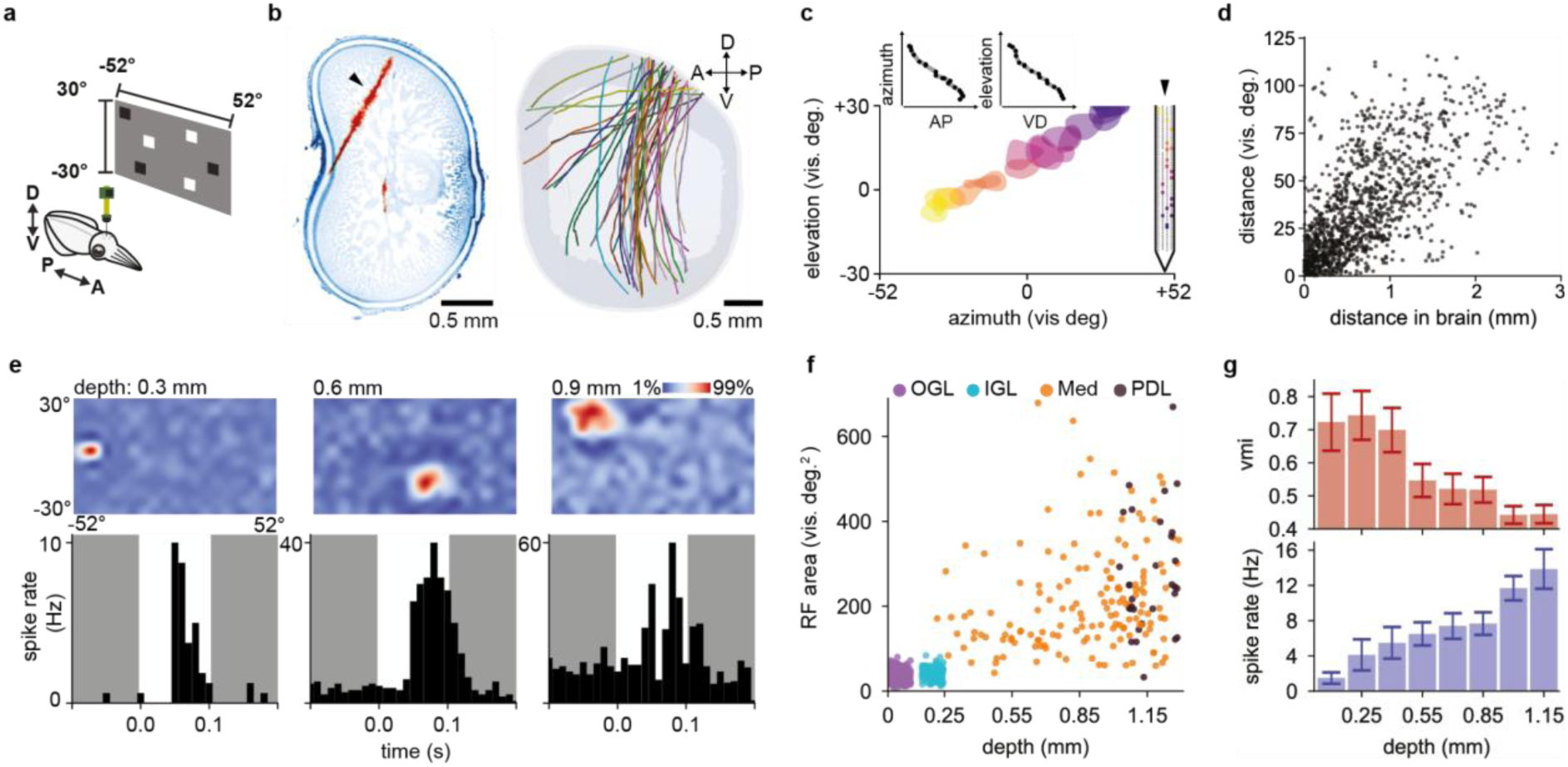
| Functional correlates of the medulla hierarchy. **a,** Schematic of sparse noise stimulation during electrophysiology recording. **b,** Left: example OL digital slice, aligned approximately sagittally to show DiI probe trace. Right: 3D OL atlas with registered probes (41 probes, 36 animals). **c,** Left: Receptive fields of a set of simultaneously recorded neurons. Right: Probe channel locations for the same set of neurons. Colour: individual neurons, ordered by probe channel. Insets: Receptive field position as a function of 3D location in the brain. **d,** Receptive field distances on the screen plotted against anatomical distances in the brain, after registration (n = 1333 neuron pairs, 36 animals, Pearson’s r = 0.675, p = 1.0 × 10^-4^ from 10,000x bootstrap resampling). **e,** Top: Receptive fields of three example neurons from three different OL depths. Bottom: corresponding spike peri-stimulus time histograms at receptive field peak locations. **f,** Distribution of receptive field sizes as a function of OL depth (n = 511 neurons, 4 animals from calcium imaging, n = 183 neurons, 30 animals from electrophysiology). **g,** Top: visual modulation index (vmi) of electrophysiologically recorded neurons with receptive fields as a function of OL depth. Bottom: corresponding baseline spike rates (n = 355 neurons, 36 animals).

Electrophysiology revealed a general increase in the range of excitatory receptive field sizes with OL depth (Fig. 4e, top row, Extended Data Fig. 11j). While neurons with relatively small receptive fields were observed throughout the OL, the distribution of receptive field sizes expanded with OL depth (0-0.6 vs 0.6-1.2mm brain depths, Mann Whitney U: 1134, p = 1.28 × 10^-3^, n = 152 neurons, 30 animals), consistent with the above imaging results from the OL cortex. Neurons in the downstream PDL showed RF sizes continuing this trend (Fig. 4f). Additionally, sparse noise visual stimuli caused local groups of electrodes to show deflections in the local field potential (LFP), allowing for an estimate of a ‘receptive field’ per electrode. These LFP receptive fields were also arranged retinotopically, and increased in size with OL depth (Extended Data Fig. 13). While superficial medulla neurons were near silent, spiking sparsely only in response to visual stimuli, deeper neurons often exhibited high levels of spontaneous activity and more temporally extended responses (Fig. 4e, bottom row). Following the trend seen in OL cortical imaging, spontaneous activity recorded electrophysiologically increased with OL depth, with visual input modulating a decreasing percent of neural activity (Fig. 4g).

### Encoding intensity and polarization contrast

To begin investigating how the medulla transforms visual information received from the OL cortex, we focused on the interaction between polarization and intensity signals. We used sparse noise stimuli in both DoLP and intensity, interleaving trials to map receptive fields across both dimensions. In the medulla, 76% of neurons with an intensity receptive field also responded to changes in polarization (45/59, n = 8 animals). Polarization receptive fields were located at similar positions to that of intensity receptive fields (azimuth: Pearson’s r = 0.86, p = 5.11 × 10^-14^, elevation: Pearson’s r = 0.79, p = 1.78 × 10^-10^), and had similar sizes (Pearson’s r = 0.61, p = 8.04 × 10^-6^). All neurons with a DoLP receptive field increased spiking activity in response to local drops in the DoLP (45/45). Many were inhibited by increases in DoLP, to a weaker extent and generally over larger areas (Fig. 5a). To test the specificity of medulla response to polarization angle, we also mapped receptive fields to DoLP stimuli following a 90 degree rotation of our polarization screen. Responsive medulla neurons showed significantly less sensitivity to this polarization angle change than seen in the OL cortex (Fig. 5a, Mann Whitney U = 22376, p = 4.17 × 10^-16^). Therefore within the medulla, upstream sensitivity to polarization angle is transformed into selectivity to polarization contrast robust to angle changes (Fig. 5b).

**Figure 5.**
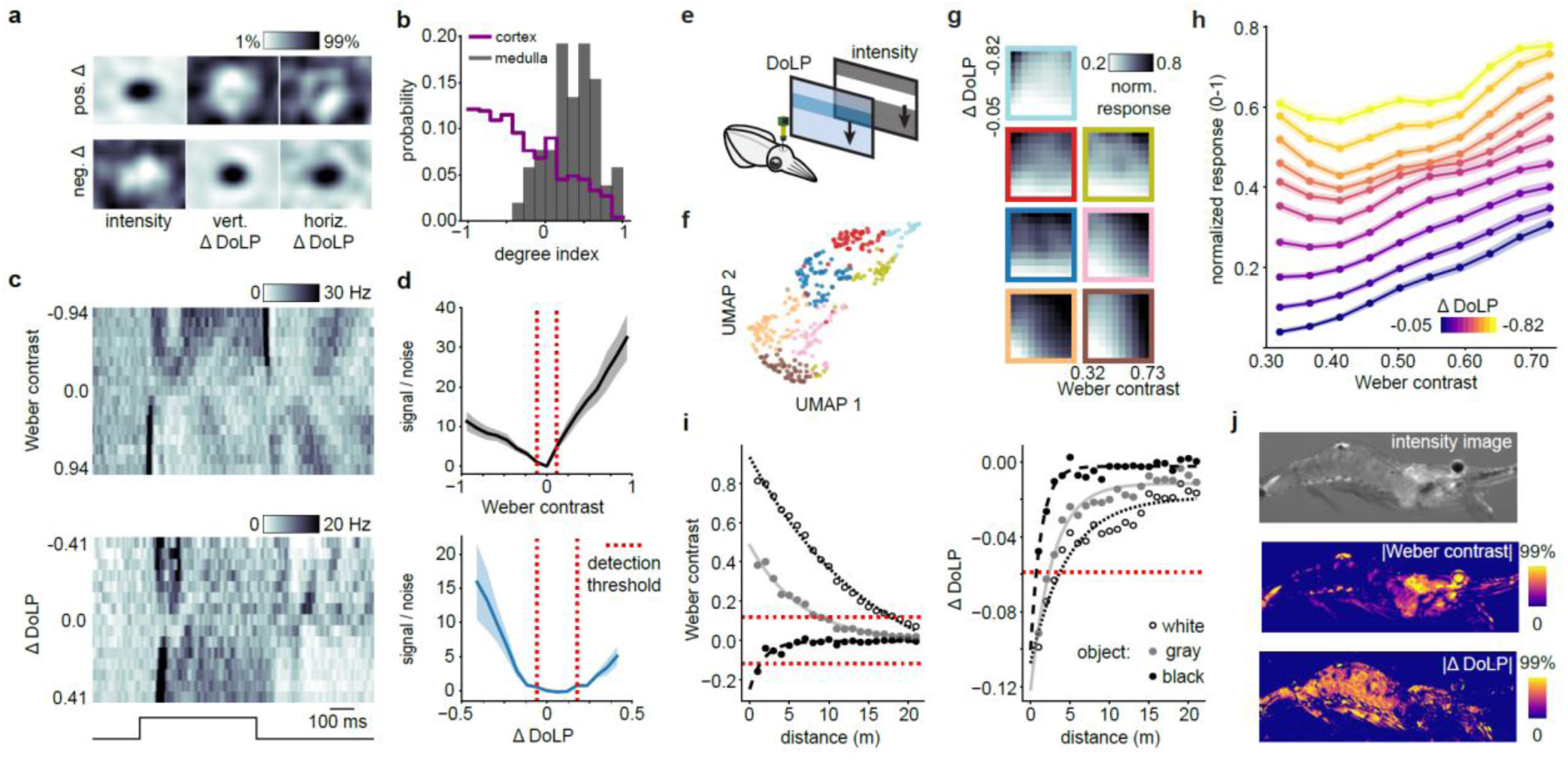
| Contrast enhancement via combining intensity and polarization signals a, Example medulla neuron RFs (60×80 visual degree crop), estimated from positive and negative sparse noise stimuli, for light intensity, ΔDoLP at vertical (vert.), and horizontal (horiz.) angles. **b**, Histogram of polarization degree index values, comparing populations of neurons in the medulla (n = 52 neurons, 8 animals) and OL cortex (pooling OGL and IGL, n = 511 neurons, 7 animals). **c**, Example trial-averaged spike rate for neurons responding to drifting bars of different Weber contrasts (top) and ΔDoLP (bottom). Bottom line marks time of stimulation. **d**, Medulla population SNR and detection thresholds for stimuli as in c. (n = 137 neurons, 6 animals for intensity stimuli, n = 116 neurons, 9 animals for ΔDoLP stimuli). **e**, Schematic of combined DoLP and intensity moving bar stimulation experiment (Methods). **f**, UMAP visualisation of medulla neural responses to combinations of intensity and ΔDoLP moving bars. Colours denote cluster identity (Leiden Clustering, n = 385 neurons, 9 animals). **g**, Cluster-averaged responses to intensity contrast + ΔDoLP stimuli. Outline colour denotes cluster identity as in f. **h**, Normalized medulla population response to intensity contrast + ΔDoLP stimuli. **i**, Left: Intensity Weber contrast and Right: ΔDoLP for black, gray, and white objects over horizontal distance from a representative dive (depth = 10m). Dots: Mean values, lines: exponential fits. Dashed lines: physiological detection thresholds. **j**, Intensity, absolute Weber contrast, and absolute ΔDoLP images of a semi-transparent shrimp imaged while diving (depth = 6 m, Methods). For the bottom two images, colour scales are set from 0 to a value of 99% of data distribution (middle: 0::0.53, bottom: 0::0.18). Values below physiological detection thresholds were set to 0.

How do squid combine light intensity and polarization degree information? We first measured contrast sensitivity to each dimension of light independently. We delivered moving drifting bars at different intensity contrast (Weber) and ΔDoLP levels on an intermediate background (gray/DoLP = 0.5, Methods), while recording from medulla neurons electrophysiologically. Most responsive neurons increased their activity monotonically to both positive and negative contrast/ΔDoLP bars, responding to the onset of lighter/negative ΔDoLP bars and the offset of darker/positive ΔDoLP bars (Fig. 5c, Extended Data Fig. 14a,b). Intensity contrast-response curves were maintained over at least a 1000x drop in light levels (Extended Data Fig. 14c). To measure an upper bound on intensity contrast and ΔDoLP sensitivity, we analysed the signal to noise ratio of the neural population at different binned contrast/ΔDoLP levels. This analysis revealed that changes in light level are detectable at both positive and negative Weber contrast bins of 11.8%. Negative ΔDoLP was detectable at 0.059, while positive ΔDoLP was only detectable at 0.177 (Fig. 5d, sequential Wilcoxon one sided test, Methods).

We next delivered combined light intensity and ΔDoLP stimuli over a range of strengths, in the form of a downward moving bar on a black/high DoLP background (Fig. 5e, Methods). Leiden clustering and UMAP visualisation of neuron responses to different contrast/ΔDoLP combinations revealed the presence of a diversity of response types (Fig. 5f). This was visible by examining neurons assigned to different clusters (Extended Data Fig. 14, as well as the cluster-averaged neural response, Fig. 5g). Some medulla neurons were more sensitive to intensity contrast, others to ΔDoLP, and many responded to the combination of both. Responses were well described by a linear additive model with a neuron specific weight for intensity contrast and ΔDoLP (R^2^ = 0.772 ± 0.008, bootstrap resampling test, 99% of neurons (381/385) had significant coefficients, Methods). Examining the distribution of intensity/ΔDoLP weights revealed that the medulla neuron population spans a continuum of sensitivity to intensity and absolute ΔDoLP increases (positive alpha, positive beta), with a small subset of neurons sensitive to ΔDoLP increases specifically at low intensity contrast (Extended Data Fig. 14, negative alpha, positive beta), and no neurons showing specific sensitivity to weak ΔDoLP stimuli (negative beta). The net result of this tuning distribution is that total medulla activity scales roughly additively with increasing light intensity or decreases in the DoLP, with a nonlinear enhancement of low intensity contrast/large ΔDoLP stimuli (Fig. 5h).

### Polarization contrast underwater

To evaluate the ecological benefits of integrating light-intensity and DoLP cues, we captured paired intensity and polarization images of calibrated targets while diving around midday (11:50-16:45) off the Okinawan coast (6-20 m depth), at locations where our squid are commonly observed (Supplementary video 4, Methods). We sampled targets of multiple contrasts, across distances and environmental conditions (cloudy-sunny, different turbidity levels, Methods). Consistent with measurements in other oceans, the background water column showed DoLP variations of 0.05-0.38 (Extended Data Fig. 15a), with foreground objects appearing as negative-DoLP silhouettes^9,32^. Both intensity contrast and ΔDoLP decayed exponentially with distance, as predicted from inherent optical properties^33–35^ and previous measurements^36^. A representative dive (depth = 10 m) shows how intensity contrast varied greatly with object brightness, falling below the detection threshold we estimated physiologically (Fig. 5d) at 1.3, 9.0, and 17.8 m for black, gray, and white objects. The ΔDoLP provided a more consistent short range signal, detectable out to 0.7, 2.3, and 3.7 m for the same targets (Fig. 5i). We analysed ten additional dives acquired under a variety of illumination and turbidity conditions. Optical properties varied markedly between dives, with white object sighting distances from 1.7-25 m using intensity contrast and 0-25 m using ΔDoLP. However, for the majority of conditions, ΔDoLP signals decayed to the detection threshold faster than those of intensity contrast (Extended Data Fig. 15d-l, 7/9 dives). This suggests that the squid’s polarization vision primarily boosts detection within the first few metres—distances where camouflage, transparency, and scattering confound intensity-based object detection despite the presence of abundant photons. While diving, we observed a particularly pronounced enhancement over intensity contrast alone for semi-transparent prey shrimp or for objects viewed against the sunlit background (Fig. 5j; Extended Data Fig. 15b).

## Discussion

Our findings reveal notable parallels between the visual system of squid and that of other organisms, suggesting common neural solutions to visual computation across evolution. Specifically, our study of the OGL and IGL revealed a diversity of functional cell types which extract distinct low-level visual features. Their retinotopy, laminar segregation, spatial coverage, convergence onto medulla targets, and the acquisition of novel features, e.g. direction selectivity, within the IGL resembles the organization of the vertebrate retina^37^, as well as the insect optic lobe^38–40^. Our data thus support Cajal’s ‘deep retina’ hypothesis^15^, suggesting that early visual processing in cephalopods, vertebrates, and insects might rely on conserved functional principles, possibly optimised for extracting reliable low-level features from complex environments^41^. The diversity of temporal response profiles we observed—such as sustained versus transient responses to intensity or polarization stimuli—also parallels known temporal coding strategies across visual systems, potentially reflecting shared computational demands for detecting and encoding dynamic visual features^30,42^. The low temporal resolution of calcium imaging, the filtering of distal electrical signals in our somatic recordings, and our use of a limited set of visual stimuli means we likely are underestimating visual response diversity in the OL cortex. The spontaneous waves of neural activity which sweep over the IGL in the absence of visual stimulation are reminiscent of spontaneous neural activity seen during vertebrate^43^ and insect^44^ development. Given the continuous and region specific growth of the cephalopod brain^18^, further studies should clarify how spontaneous activity shapes the development of network connectivity or its maintenance. Moreover, given the association we observed between spontaneous activity waves and behavioural state changes (e.g., breathing and twitching), future studies could explore how arousal or sleep-like states^31^ modulate visual processing in cephalopods, analogous to state-dependent modulation described in vertebrates^45^ and insects^46^.

The OL medulla’s tree-like anatomical organization and receptive field structure suggest a hierarchical organization of visual information processing in the cephalopod brain. This also resembles visual hierarchies across vertebrates^41,47^ and insects^48^. While receptive fields generally increased with OL depth, we also recorded deep medulla neurons with small receptive fields. This hints at the presence of OL ‘skip connections’, a strategy for preserving high-resolution details in deep neural networks^49^. Consistent with this possibility, we filled some neurons which possessed long axons connecting superficial and deeper OL over at least 300μm (Extended Data Fig. 8b). Although our data are consistent with a predominantly feed-forward architecture, anatomical studies have profiled the presence of dense feedback from central brain regions to the OL, intra-laminar recurrent loops, and commissural fibres linking the two optic lobes^13^ (Extended Data Fig. 9c). In mammalian vision, feedback connections are thought to modulate attention^50^, predictive coding^51^, and behavioural states^52^. Determining whether analogous top-down processes operate in cephalopods will require custom behavioural paradigms or neural manipulations capable of probing attention-like states or predictive processing in these animals. Of equal importance is describing whether and how medulla neurons integrate the diversity of signals from the OL cortex to represent the kind of higher order features observed in the mammalian cortex, e.g. visual textures used for camouflage behaviour^53^.

Our results detail the transformation of retinal-level polarization sensitivity into a format which provides behavioural advantages to the squid. Inputs from photoreceptors tuned to specific polarization angles are combined within the superficial OL to produce neurons with a range of angle specificities. Downstream medulla neurons become equivariant to polarization angle, acquiring instead sensitivity to local decreases in the DoLP, which they integrate additively with luminance differences into a general contrast signal. While this computation explains behavioural findings that polarization cues help many species detect objects when intensity contrast alone is unreliable^3–10,54,55^, the underlying circuit mechanisms in cephalopods—whether selective synaptic convergence, inhibitory interactions, or both—remain unclear, and await further investigation by connectomics or targeted electrophysiology.

Our fieldwork measurements demonstrate that intensity- and polarization-based contrasts in underwater scenes are both strongly, yet independently, shaped by lighting conditions and water turbidity: under some circumstances intensity contrast provides the dominant signal, whereas in others polarization cues prevail. Our measurements also demonstrate how some prey animals, such as the semi-transparent glass shrimp, can minimise their intensity-based contrast against the background water column, yet their polarization silhouette remains conspicuous even at short viewing distances when photon flux is plentiful. Possessing independent, highly sensitive intensity and polarization contrast channels therefore enables cephalopods to detect relevant visual cues over a wider range of behavioural and environmental conditions^56^.

Recent work in zebrafish suggests that vertebrate chromatic sensitivity and the neural circuits mediating it may have evolved to use the rapid filtering of ultraviolet and red light to assist in basic functions like contrast enhancement and judging distances underwater, with true colour vision emerging from these more basic computations^57^. Our data suggests that the cephalopod visual system employs polarization sensitivity to support similar functions, albeit utilizing a distinct dimension of light^58^. The molluscan ancestors of cephalopods possessed multiple opsins, suggesting that modern cephalopods underwent a reduction of opsin types, perhaps to prevent ambiguous signals generated by photoreceptors sensitive to both wavelength and polarization properties of light^59^. Testing this hypothesis would require detailed comparative genomic and physiological analyses across extant species, potentially illuminating evolutionary trade-offs between polarization and colour sensitivity. Notably, many crustaceans possess high-resolution polarization vision and only a single opsin, or segregate their neural circuits for polarization and wavelength discrimination^60^. Insects use polarization sensitivity very differently than cephalopods, but they too mostly segregate polarization and colour visual circuits functionally^61,62^(but see^63,64^), as well as morphologically removing polarization sensitivity in colour pathways through rhabdomere twisting^65^.

More generally, our work reveals points of convergence between cephalopod vision and that of other neural systems. The vast evolutionary distances to other biological systems suggest that these convergent features reflect shared functional constraints on the emergence of complex visual systems^66,67^. The common initial processing of visual scenes using a set of distinct feature detectors may reflect the principle of efficient coding^68–71^. Visual hierarchies possibly form repeatedly due to the compositional nature of natural images^72,73^. And polarization and colour vision may have both evolved due to the common need to boost contrast in underwater environments^3,58,59,74^. Together, our observations highlight the potential for cephalopods to serve as models for studying fundamental neural computations. The many parallels between the cephalopod OL and other visual systems prompt further investigation into whether similar synaptic and circuit-level mechanisms underlie these convergent outcomes. Future experiments adapting genetic techniques such as cell-type-specific labeling, optogenetics, or CRISPR-based manipulations from established model organisms to cephalopods could significantly facilitate these investigations.

## Figures and Legends

**Extended Data Figure 1.**
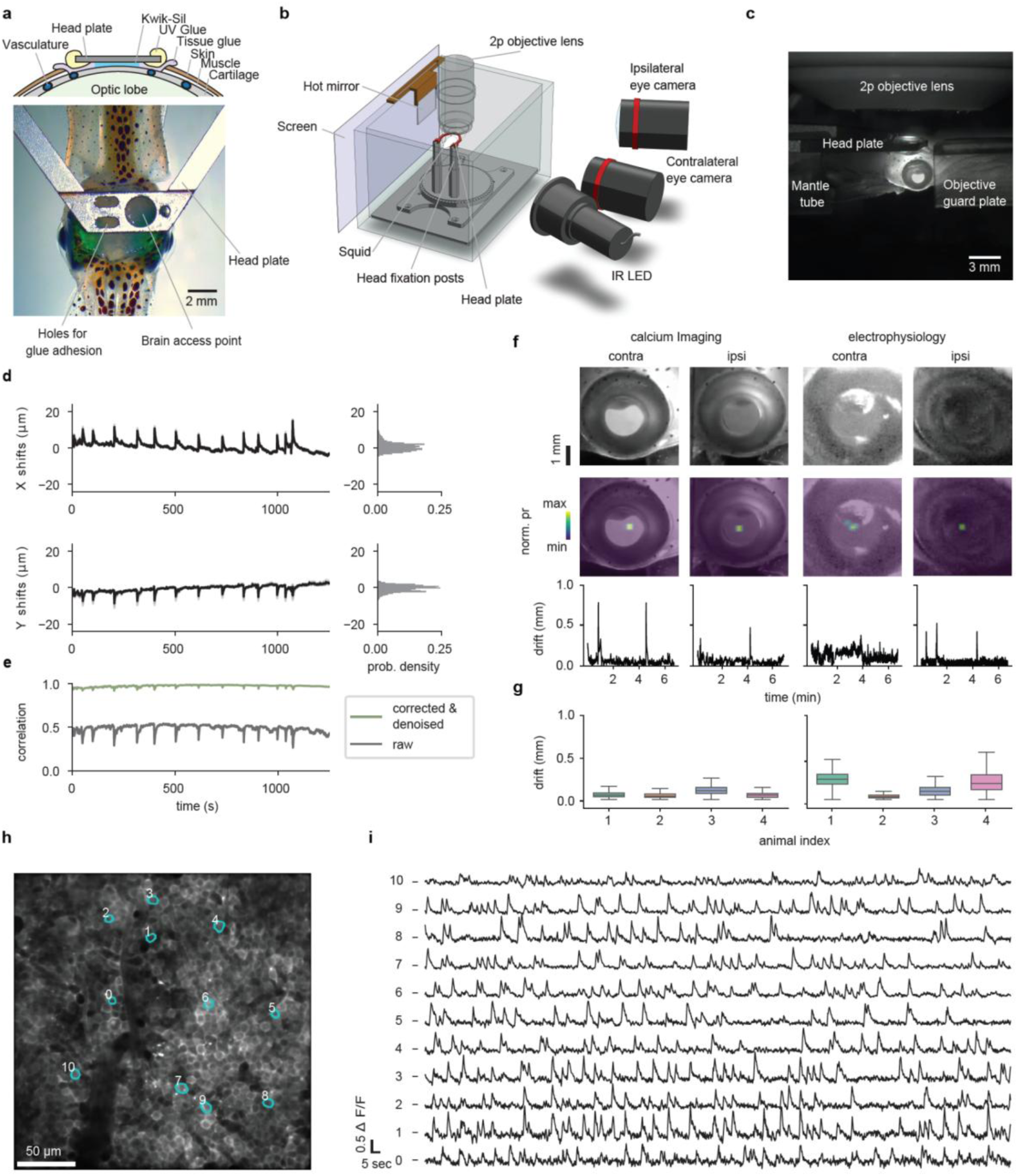
| In vivo calcium imaging in awake squid. **a**, (Bottom) Photograph of a head-fixed squid. (Top) Schematic of the surgical and gluing procedure. Although squids lack rigid skeletal structures, their brains are encased in a thin sheet of cartilage, to which a metal head plate was attached. Imaging/probe insertion proceeds through the circular hole in head-fixation plate. Smaller holes are for increasing adhesion. **b**, Schematic of head fixed recording setup. **c**, Squid viewed from the side (contralateral eye camera). **d**, Motion drift trace from a representative in vivo calcium imaging session in the OGL, quantified using the motion correction algorithm NoRMCorre. Strong jetting behaviour by the squid occasionally appears as spikes in the drift trace. **e**, Frame-wise correlation between each image frame and a reference template. The black line shows correlation values for the raw movie prior to correction; the green line shows values after non-rigid motion correction and denoising, demonstrating the stability of the recording over a 20-minute session. **f**, Eye movement tracking during Ca^2+^ imaging (left) and electrophysiology recordings (right). Top row: cropped images of eyes ipsi- and contra-lateral to visual stimulation. Second row: probability density of tracked eye position. Third row: example drift traces (deviation from mean eye position). **g,** Distribution of eye drifts for the ipsilateral eye during Ca^2+^ imaging (left) and electrophysiology recordings (right) over multiple animals, 6.8 minutes per recording. **h–i**, Representative calcium transients recorded from OGL neurons during presentation of sparse noise stimuli. h, Image of the imaging field with labeled regions of interest (ROIs). i, ΔF/F traces corresponding to the ROIs shown in d.

**Extended Data Figure 2.**
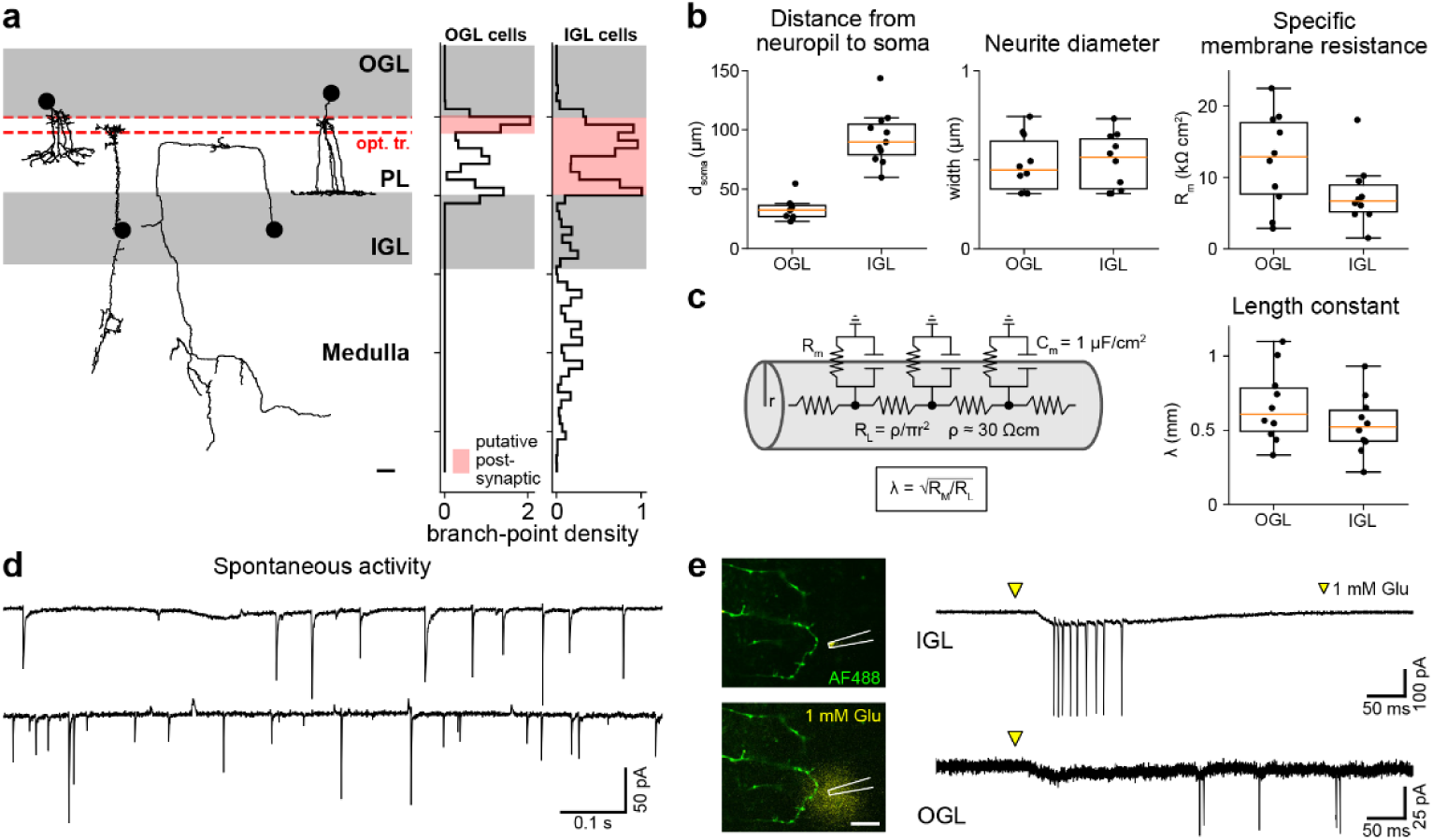
| Signal propagation in OL cortical neurons. **a,** Left: Example morphological reconstructions of neurons in the OGL (n=10) and IGL (n=10). Traces are aligned to the tangent plane of the OL surface and normalized to the thickness of the plexiform layer (PL, scale bar = 0.2 * PL width, average PL width=54.66 ± 0.75 µm). Nerve endings of the optic tract are found in the area denoted by the red dashed line. Right: Distribution of branch-points as a function of depth. Red shaded regions denote areas of putative postsynaptic terminals. **b**, Left: Distance, measured along neuronal processes (n=10 OGL, 10 IGL neurons), from branch-points of putative postsynaptic terminals to the soma (OGL: d_soma_ = 33.09 ± 3.05 µm, IGL: d_soma_ = 93.30 ± 6.51 µm). Middle: Diameter of the primary neurite extending out of the soma (OGL: 0.48 ± 0.05 µm, IGL: 0.49 ± 0.05 µm). Right: Specific membrane resistance of OGL and IGL neurons (OGL: R_m_ = 12.389 ± 1.993 kΩ cm^2^, IGL: R_m_ = 7.578 ± 1.329 kΩ cm^2^). **c**, Signal propagation along neurites can be approximated by a cylindrical cable with longitudinal resistance R_L_ and membrane resistance R_M_. R_L_ was estimated from the axoplasmic resistivity (ρ ≈ 30 Ω cm^75,76^) and the radius of the neurite (Methods). The specific membrane capacitance was taken^77^ to be 1 µF/cm^2^. The rate of decay along the cable is given by the length constant λ (OGL: λ = 0.66 ± 0.07 mm, IGL: λ = 0.54 ± 0.06 mm). **d**, Example traces of spontaneous activity recorded in two IGL neurons during voltage clamp. Spontaneous EPSCs and IPSCs are detected at the soma. **e**, IGL and OGL neurons were stimulated by focal pressure application of 1 mM L-Glutamate (+ 25 µM Alexa Fluor 594) next to putative postsynaptic terminals via a glass pipette. Glutamate puff often induced action potential firing, either directly (top trace) or apparently polysynaptically (bottom trace), which propagated back to the soma. Scale bar = 10 µm.

**Extended Data Figure 3.**
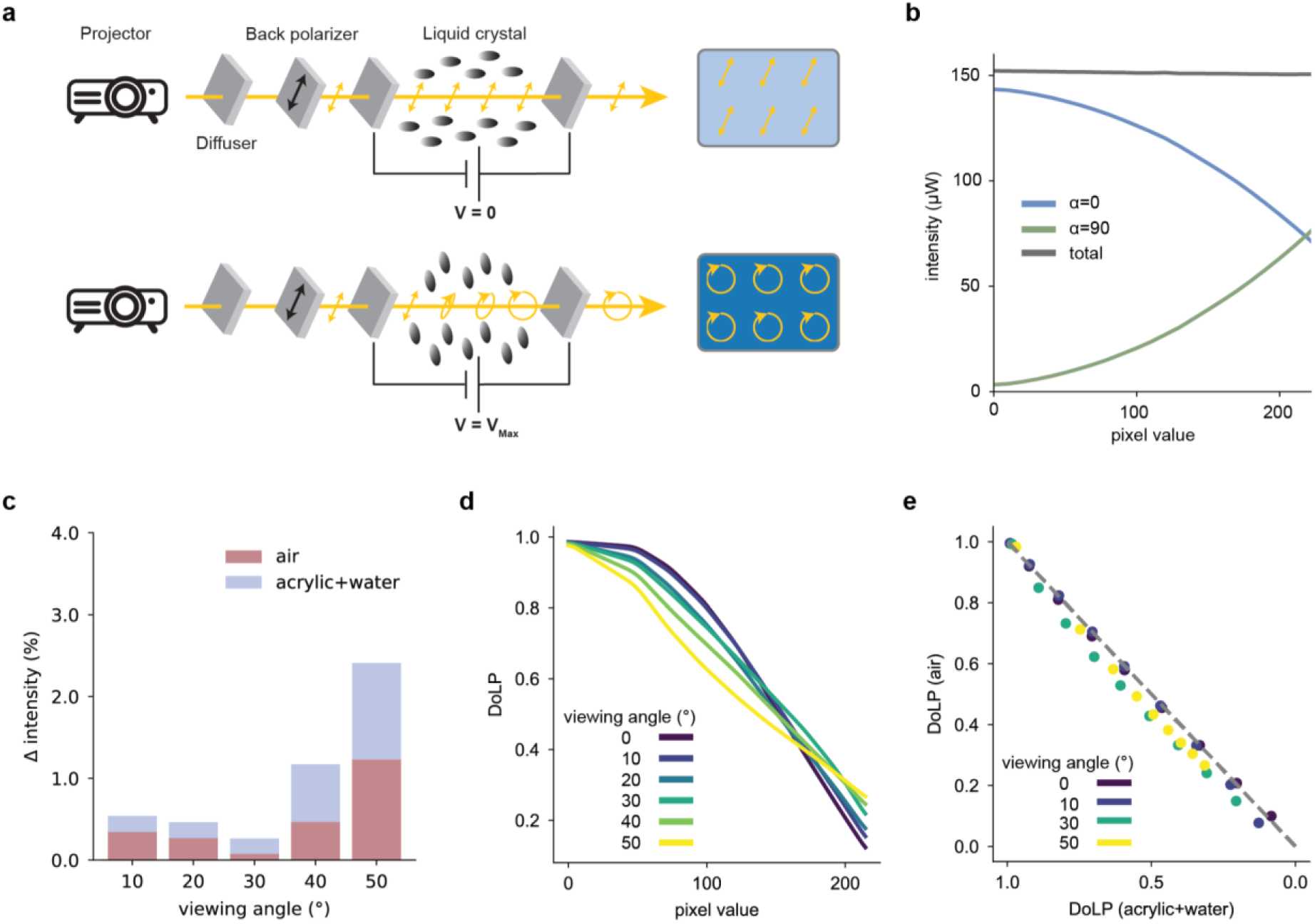
| Screen calibration. **a,** Schematic of device used to deliver light intensity and polarization visual stimuli^27^. **b,** Calibration of light intensity levels as a function of LCD pixel value, measured with a photodiode, through a linear polarization filter at horizontal (α = 0) and vertical (α = 90) angles, as well as overall (total). **c.** Change in light intensity between polarized (LCD pixel value 0) and unpolarized (LCD pixel value 220) light as a function of viewing angle, with air and viewed through an experimental tank filled with seawater. **d.** DoLP as a function of LCD pixel value, under different viewing angles. **e.** DoLP values between air and the experimental tank filled with seawater as mediums, under different viewing angles.

**Extended Data Figure 4.**
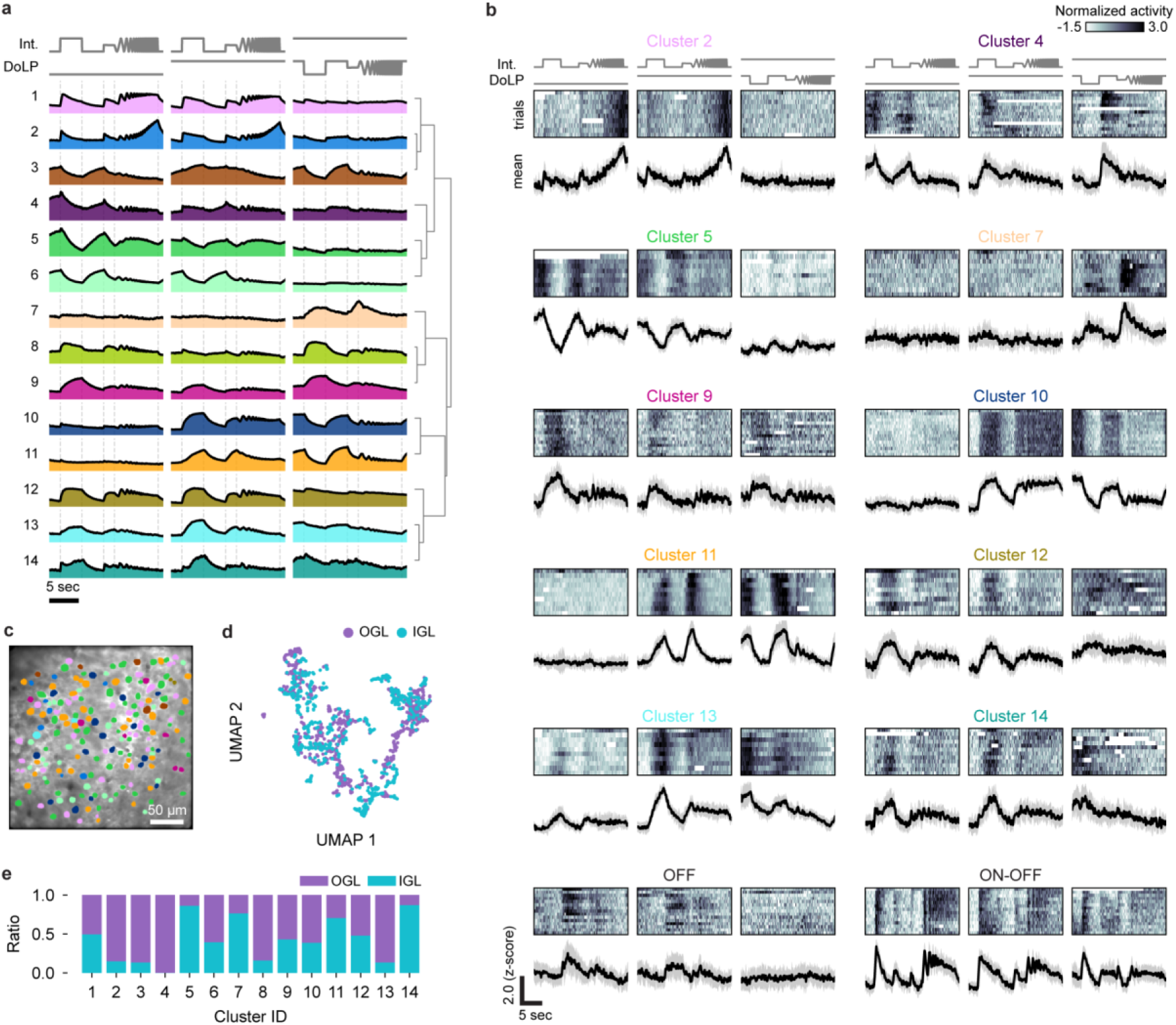
| Functional clustering of neurons in the OL cortex. **a**, Mean Ca^2+^ response of 14 functionally distinct clusters (also see Fig. 1). Dendrogram represents similarity relationships between clusters. These clusters are the result of manual merging of similar functional responses, following algorithmic clustering (Methods). **b,** Example single-neuron responses from each cluster, along with two cells (“OFF” and “ON-OFF”) with distinct functional properties which did not form clusters due to sparsity. **c,** Spatial distribution of functional cell types in a representative IGL recording (n = 191 neurons). **d,** UMAP embedding of all neurons, coloured by region identity. **e,** Ratio of OGL and IGL neuron numbers in each cluster.

**Extended Data Figure 5.**
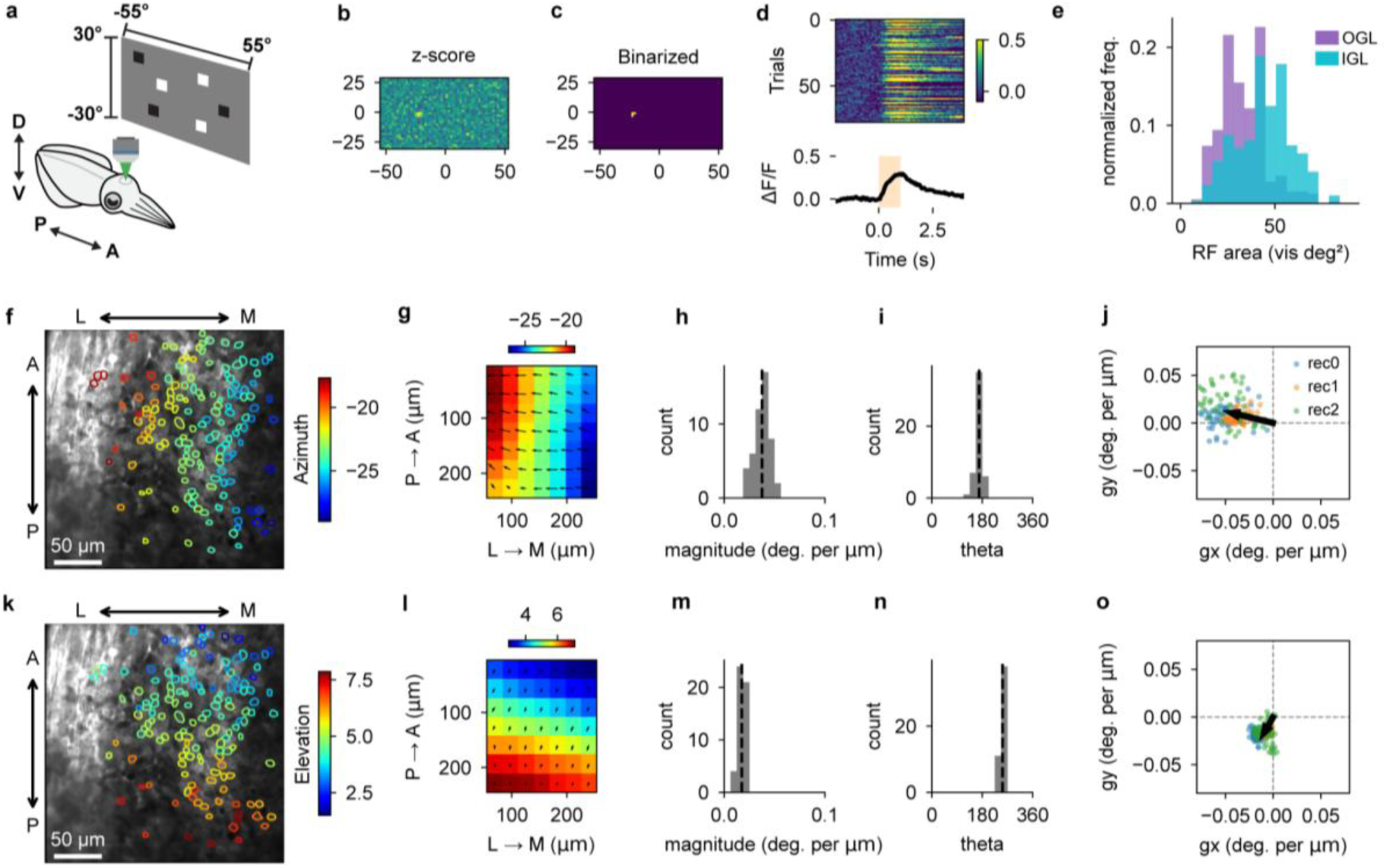
| Receptive field mapping in the superficial OL. **a**, Schematic of the visual stimulus and recording setup. **b–d**, Receptive field (RF) of an example neuron in the OGL. b, Pixel-wise response amplitude shown as z-scores. c, Binarized image used to define the RF. d, Event-triggered average of calcium signals (bottom) from trials with a stimulus in the RF (stimulation timing highlighted in beige; top panel shows single-trial activity heatmap). **e**, Distribution of RF area for OGL neurons (n = 394 neurons, 3 animals) and IGL neurons (n = 297 neurons, 2 animals). IGL RF size was significantly larger than OGL (p = 1.03 x 10^-24^, Welch’s t-test). **f**, Azimuth map of RF centres in one representative recording in the OGL. Each cell is colour-coded by its azimuthal RF position. **g**, Spatial gradients of azimuthal RF positions were estimated by fitting a vector field. **h**, Distribution of gradient magnitudes (degrees per µm). **i**, Distribution of gradient directions (theta, in degrees). **j**, Scatter plot of gradient vectors (gx, gy) from three recordings. Colours indicate datasets; black arrow denotes the mean gradient. **k–o**, Same analyses as in f–j, for elevation values of RF centres.

**Extended Data Figure 6.**
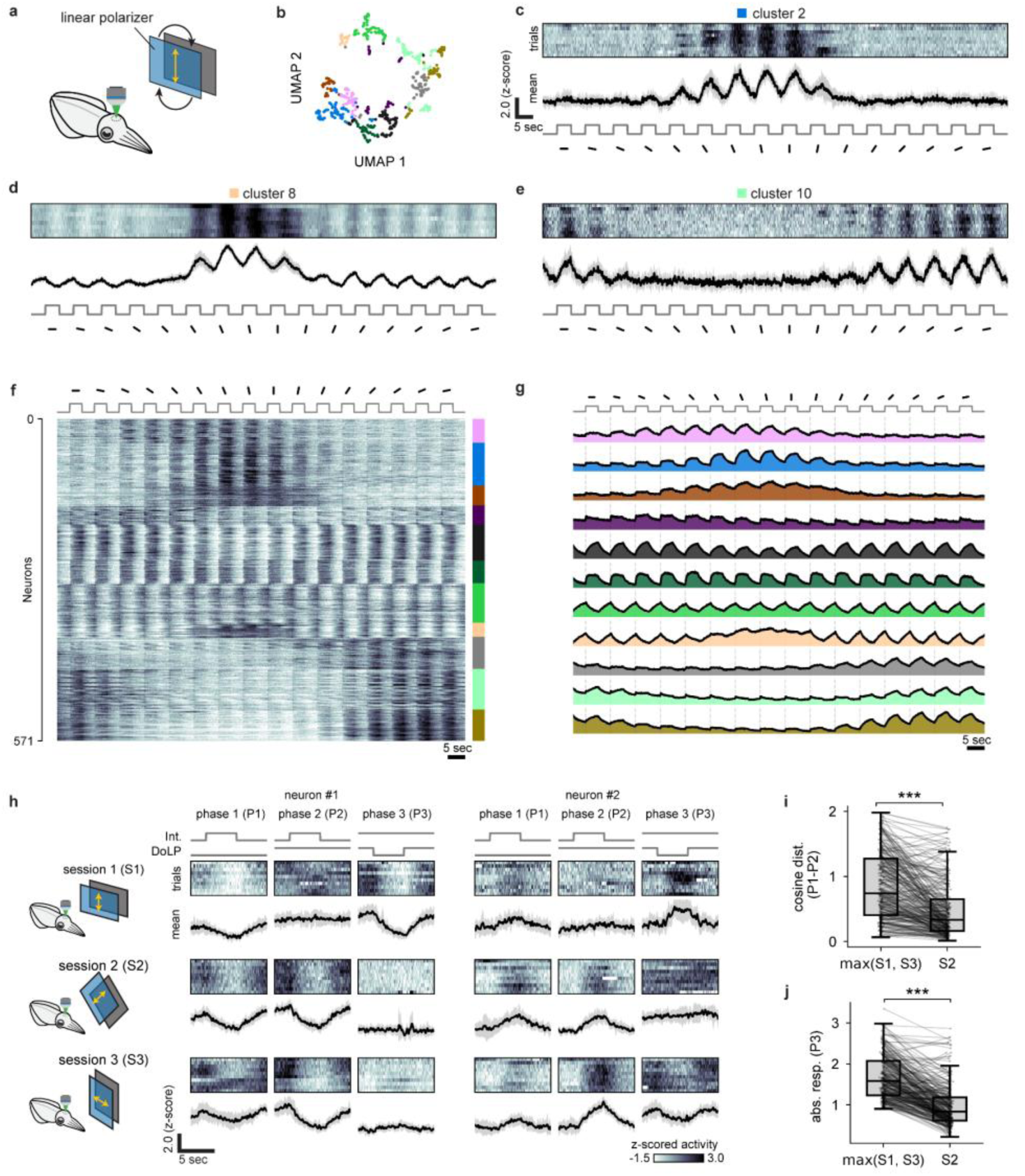
| Two channels for processing polarized light. **a**, Schematic of the linear polarizer rotation experiment. Full field stimuli were presented at 16 different polarization angles (Methods). **b**, UMAP embedding of all neurons’ responses, coloured by cluster identity. **c-e**, Example Ca2+ responses from single neurons, taken from three different clusters. **f**, Pooled dataset of OGL (n = 497 neurons, 3 animals) and IGL (n = 74 neurons, 2 animals). Colours: cluster identity. **g**, Cluster averaged Ca2+ responses. **h**, Schematic and example neural responses to vertical, horizontal and 45-degree stimulation. **i**, Neural responses to polarized vs unpolarized intensity stimulation (P1-P2) are more distinct at vertical/horizontal angles (S1 or S3), than at 45-degree stimulation (S2, paired t-test, p = 7.47e-31, n = 264 neurons from 3 animals). **j**, Neural responses to DoLP stimulation (P3) are stronger at vertical/horizontal angles (S1 or S3), than at 45-degree stimulation (S2, paired t-test, p = 3.38e-55, n = 264 neurons from 3 animals).

**Extended Data Figure 7.**
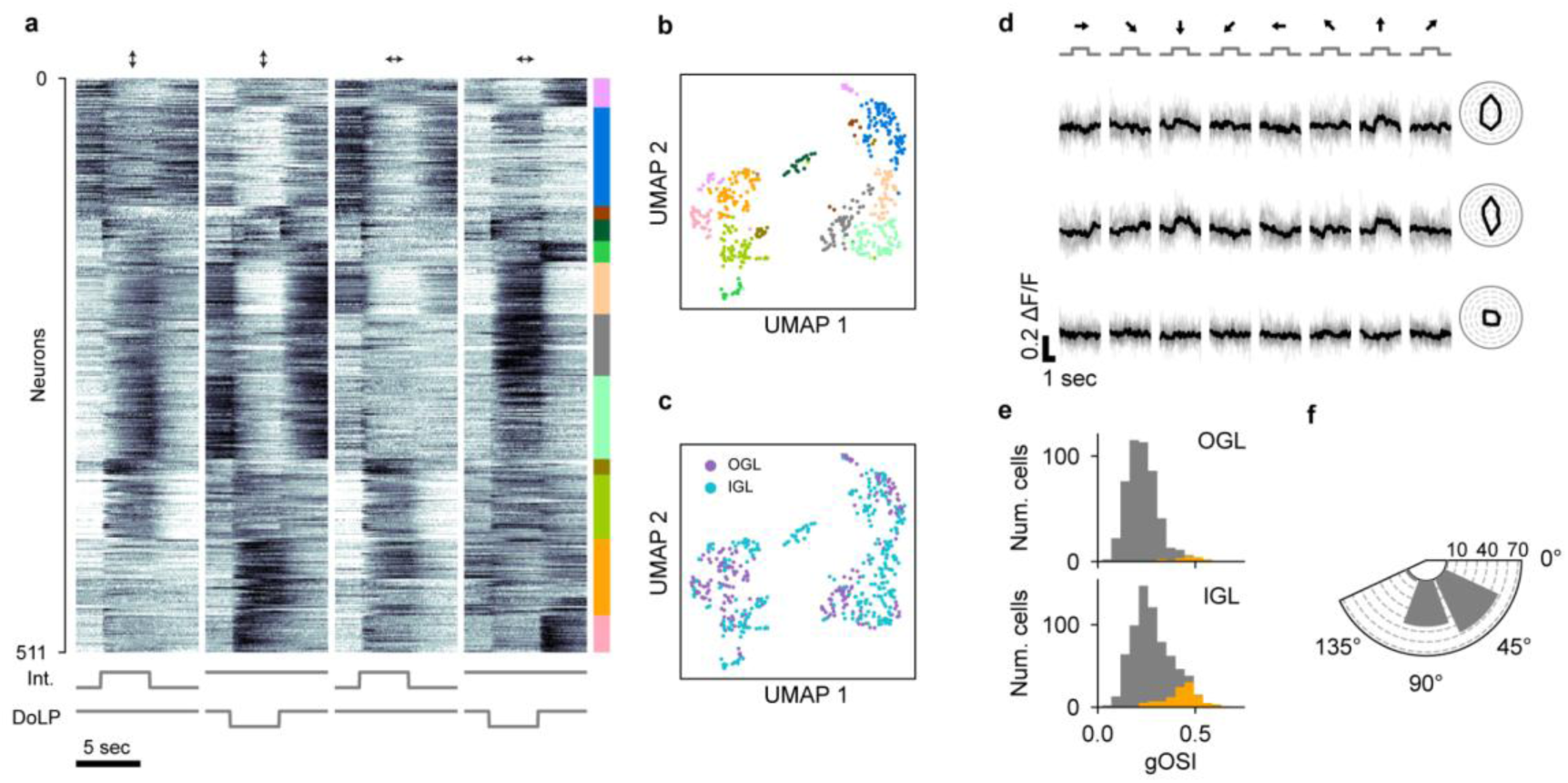
| Polarization angle tuning and orientation-selective neuron in the OGL/IGL. **a**, Clustering of Ca^2+^ responses to intensity/ΔDoLP stimuli with vertical (first two columns) and horizontal (last two columns) screen orientations. **b**, UMAP embedding of all neurons, coloured by cluster identity. **c**, UMAP embedding of all neurons, coloured by region identity. **d**, An example orientation-selective (OS) neuron in the IGL. Top row: intensity grating, unpolarized background. Middle row: intensity grating, vertically polarized background. Bottom row: DoLP grating, constant-intensity background. **e**, Distribution of gOSI values for neurons in the OGL (top) and IGL (bottom). Orange bars indicate OS neurons with significant orientation tuning (n = 14/491, 2.8% in OGL, n = 109/695, 15.6% in IGL). **f**, Distribution of preferred orientations among OS cells in the IGL (n = 109).

**Extended Data Figure 8.**
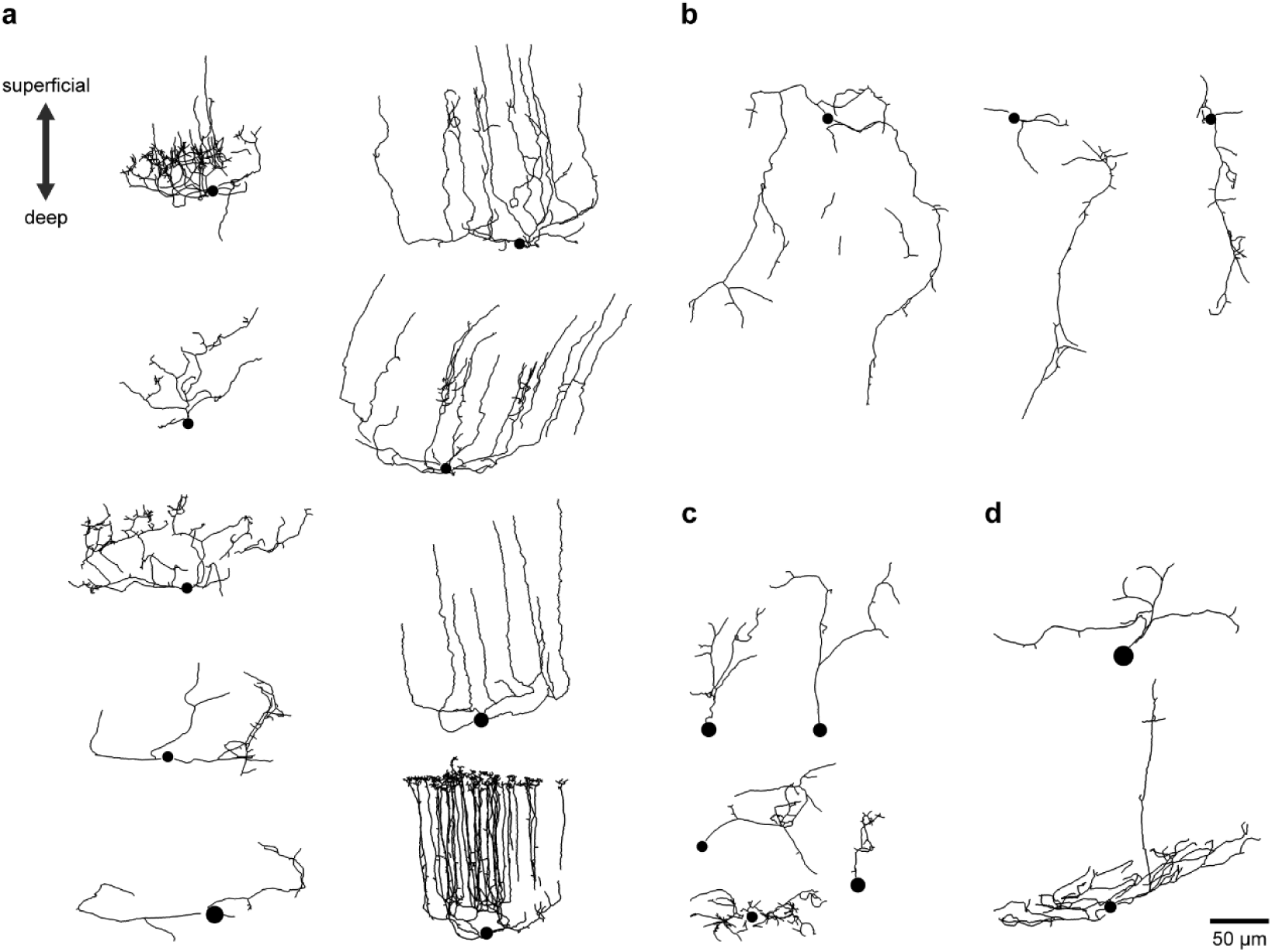
| Single cell neural morphology within the medulla. Single neuron morphology revealed by intracellular dye filling and tracing of processes. Neurons are rotated to a common superficial-deep OL axis. Spheres indicate soma location. **a**, Neurons with processes oriented towards the OL surface. **b**, Neurons with processes oriented away from the OL surface. **c**, Neurons with processes restricted to the local cell island. **d**, Miscellaneous/uncategorized neurons.

**Extended Data Figure 9.**
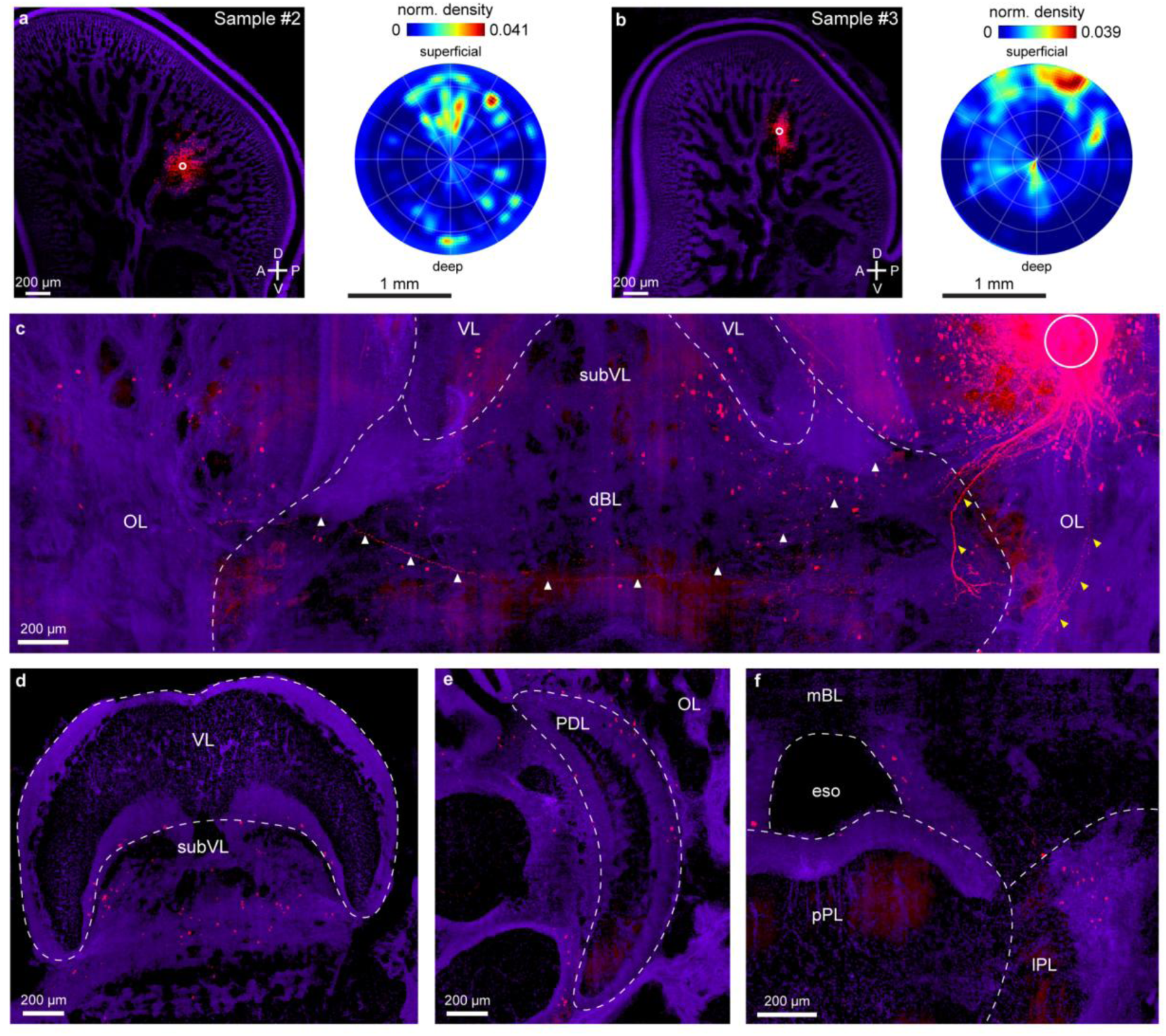
| Circuit tracing of the OL medulla using CM-DiI. **a**, Left: Tracing result from Sample #2 showing CM-DiI (red) labelling in the OL medulla, overlaid on nuclear staining (purple). Sagittal view, maximum intensity projection of a 50-μm thick volume. Circle: injection site. Right: 3D normalized density of DiI-labelled somata mapped onto a 2D polar plot (Methods). The injection site is at the centre, and the vertical axis represents the superficial-to-deep OL axis. **b**, Tracing result from Sample #3. **c**, Coronal section from Sample #2, maximum intensity projection of a 500-μm thick volume. The injection site is marked by a circle. White arrowheads indicate contralaterally projecting OL axons, while the yellow arrowheads mark axons connecting to the lateral basal lobe and peduncle lobe. Soma-localized CM-DiI labelling is observed in neurons of the contralateral OL, as well as in the subvertical lobe (subVL) and dorsal basal lobe (dBL). **d-f**, Maximum intensity projection of a 300-μm thick volume, showing detailed views of **d**: vertical lobe (VL) and subVL, **e**: peduncle lobe (PDL), and **f**: median basal lobe (mBL), posterior pedal lobe (pPL), and lateral pedal lobe (IPL) regions. eso: esophagus.

**Extended Data Figure 10.**
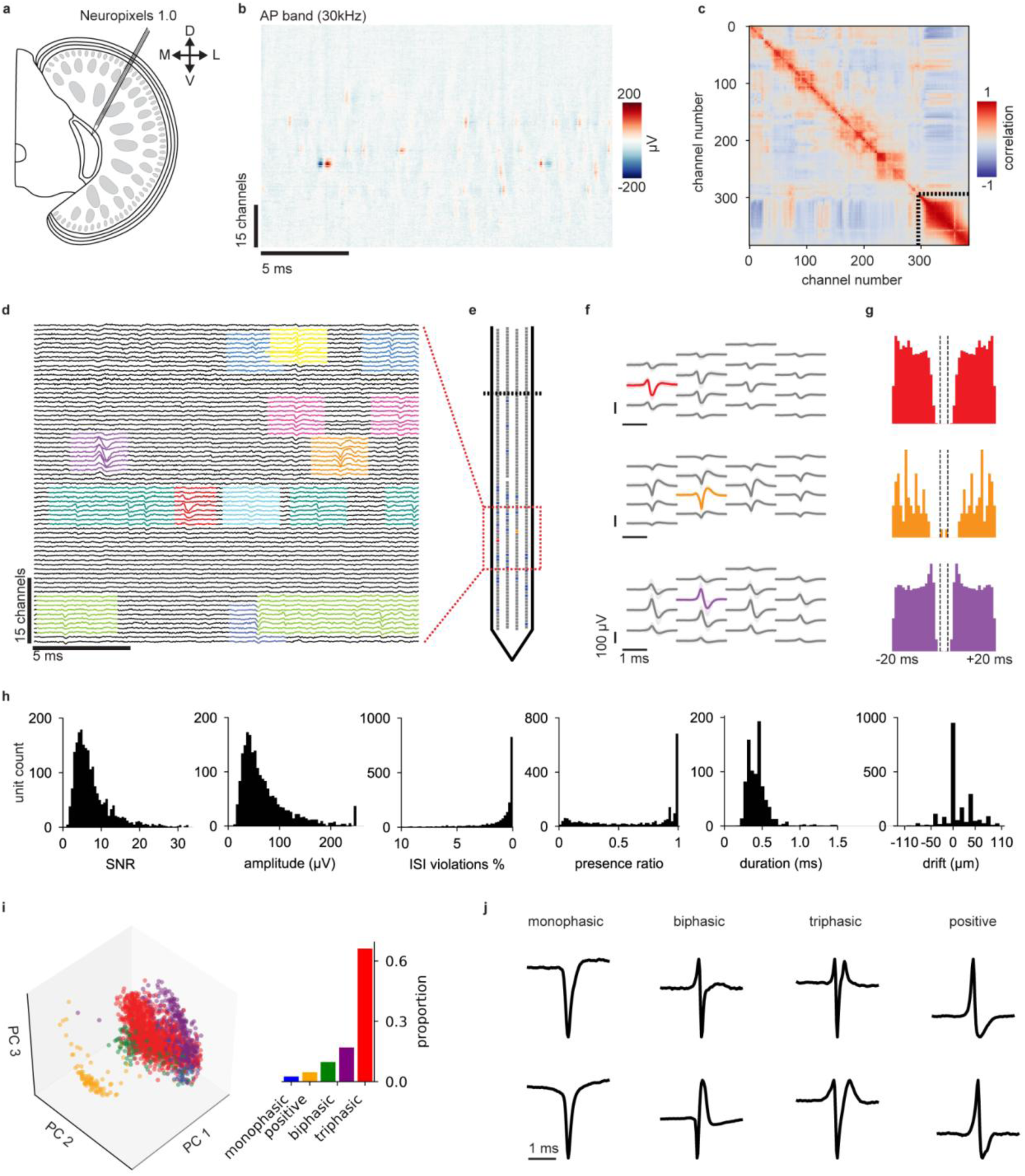
| In vivo electrophysiology data pipeline. **a,** Schematic of Neuropixels probe insertion into the OL. **b**, Example high pass filtered LFP (Neuropixels action potential band) recording from 384 channels of the probe, local deflections represent unit activity. **c**, Correlation matrix of low pass filtered LFP (Neuropixels LFP band). Stippled lines denote channels located outside of the brain, detectable through their distinct correlation structure. **d**, Example waveforms assigned to single units (colours) following spike sorting and manual curation. **e**, Location of units from d on the Neuropixels probe. Stippled line denotes the brain boundary. **f**, Example simultaneously recorded single unit waveforms across local groups of channels. Colours match waveforms in d. **g**, Inter-spike interval distributions for units in f (n= 16743, 2293, 48727 spikes for each unit respectively). **h**, Histograms showing distribution of spike sorting quality statistics (n= 2142 units, 47 animals). **i**, Example waveform shapes. **j**, (Left) Proportion of neurons belonging to each spike waveform class. (Right) Principal component projection of waveforms reveals separation of positive class.

**Extended Data Figure 11.**
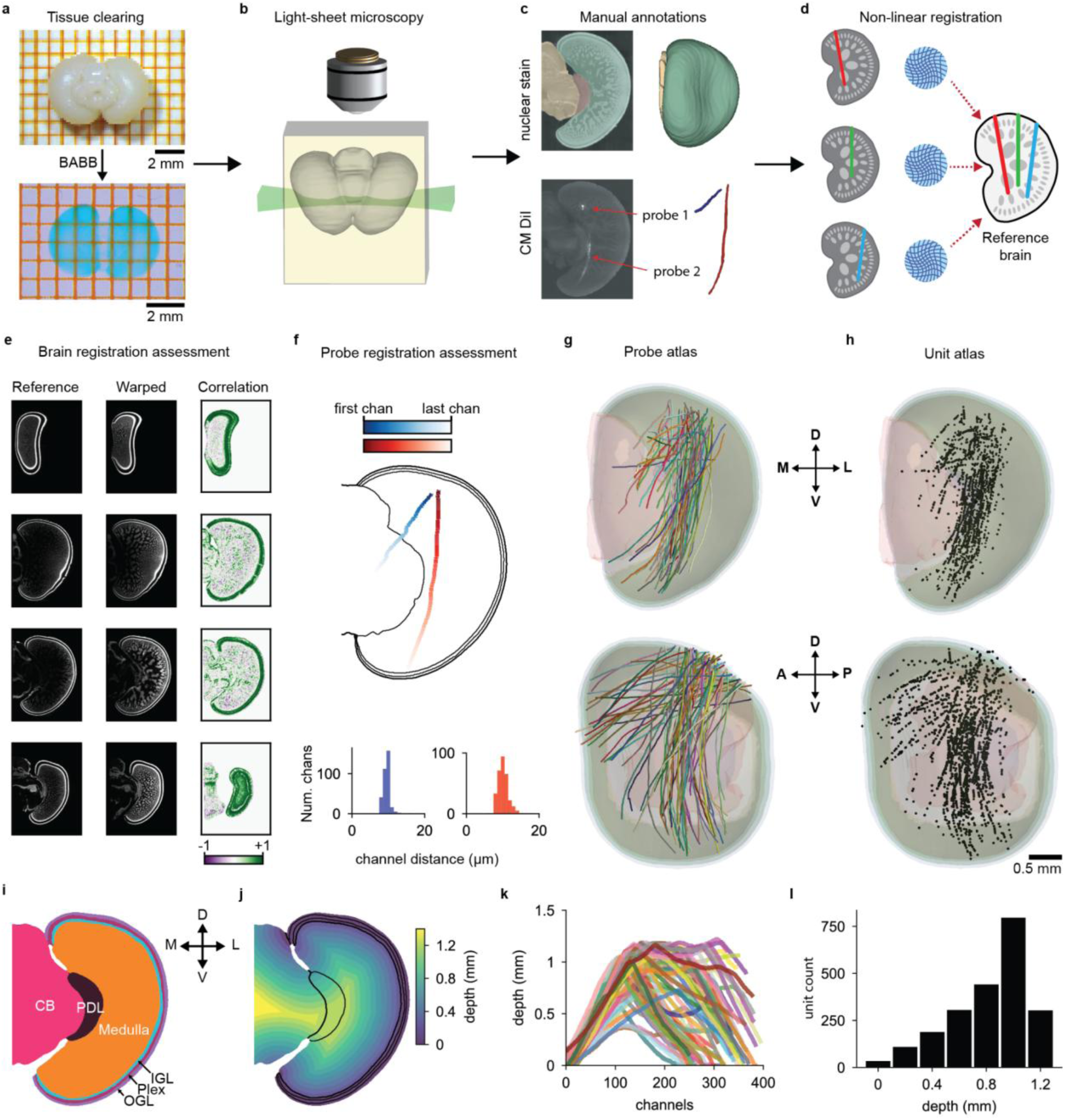
| Squid brain atlas and 3D probe registration pipeline. **a-d,** Neuropixels probe registration pipeline. **e,** Warping quality assessment through cross correlation. **f**, Skeletonization and warping quality assessment through channel sequence, position and inter-channel distance evaluation. **g**, Resulting brain atlas with registered probes (71 probes, 47 animals). **h**, Resulting brain atlas with unit positions for all probes (n = 2142 units). **i**, Brain region segmentation of reference brain. **j**, OL depth map calculated from a distance transform from OL exterior. **k**, OL depths traversed by all probes. **l**, Number of recorded units as a function of depth.

**Extended Data Figure 12.**
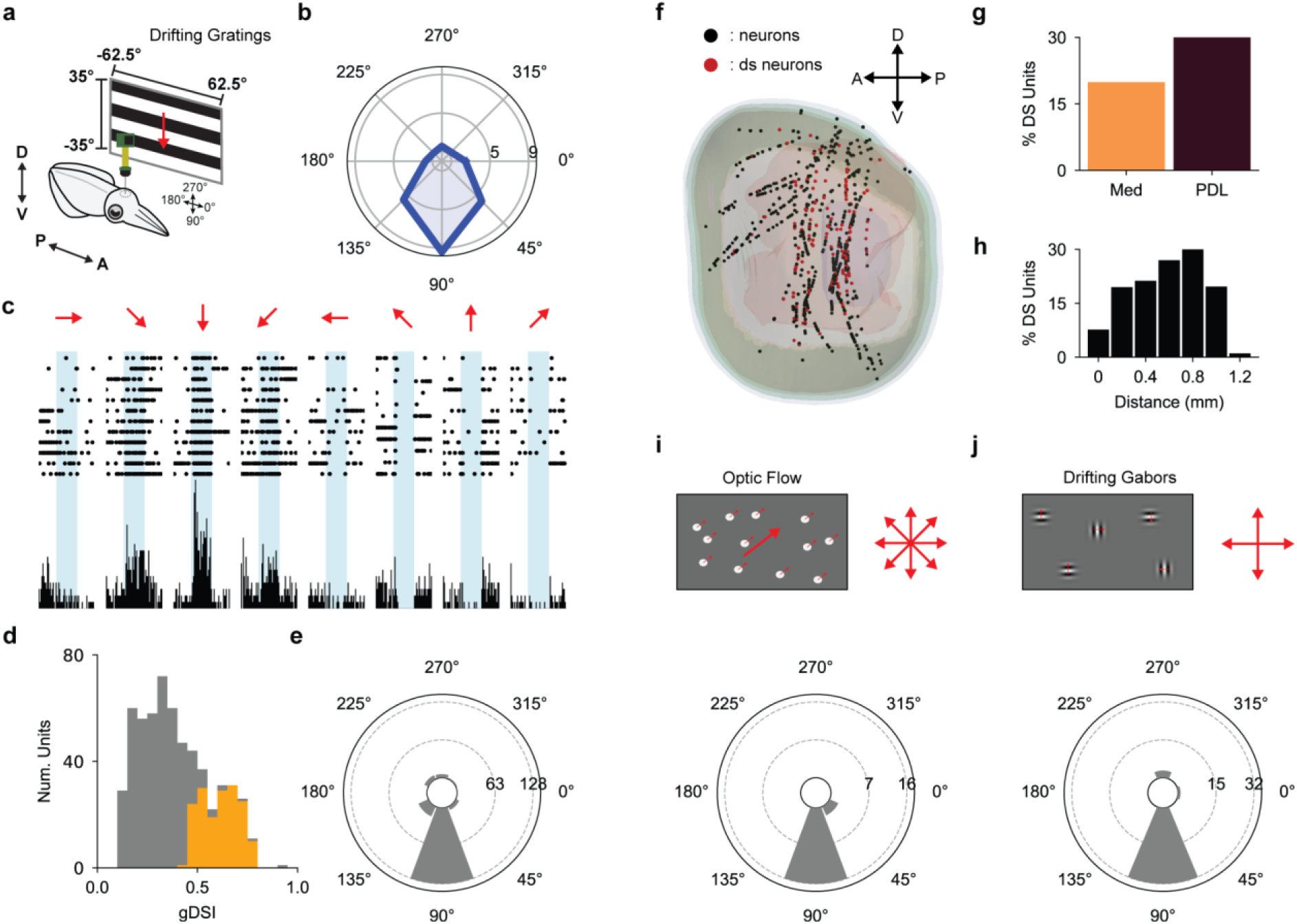
| Direction selectivity within the medulla. **a,** Schematic of drifting grating experiment. Stimuli span directions, spatial (0.024, 0.06, 0.18, 0.26, 0.84, 1.2 cycles/degree) and temporal frequencies (0, 1.5, 3, 7 Hz). **b,** Firing rate distribution of an example neuron’s direction selectivity at preferred spatial and temporal frequencies (Methods).. **c,** Raster plot (top) and PTSH (bottom) of neuron in b. **d,** Distribution of gDSI values for neurons recorded with NP, orange bars indicate statistically significant gDSI values (22% direction selective neurons, n = 169 visually responsive neurons, 18 animals). **e,** Direction selectivity distribution of all DS-selective neurons in d. **f,** Sagittal view of OL atlas showing location of all neurons recorded during drifting gratings experiments (black) and DS-selective neurons (red). **g,** Percentage of DS-selective neurons in the OL medulla and the peduncle lobe. **h,** Percentage of DS-selective neurons as a function of OL depth. **i,** In a subset of experiments, optic flow stimuli were presented to the animal (top), with a similar distribution of DS-selective neurons (bottom) (7% direction selective neurons, n = 19 visually responsive neurons, 5 animals). **j,** In a subset of experiments, drifting local Gabors were presented to the animal (top), with a similar distribution of DS-selective neurons (bottom) (9% direction selective neurons, n = 38 visually responsive neurons, 9 animals).

**Extended Data Figure 13.**
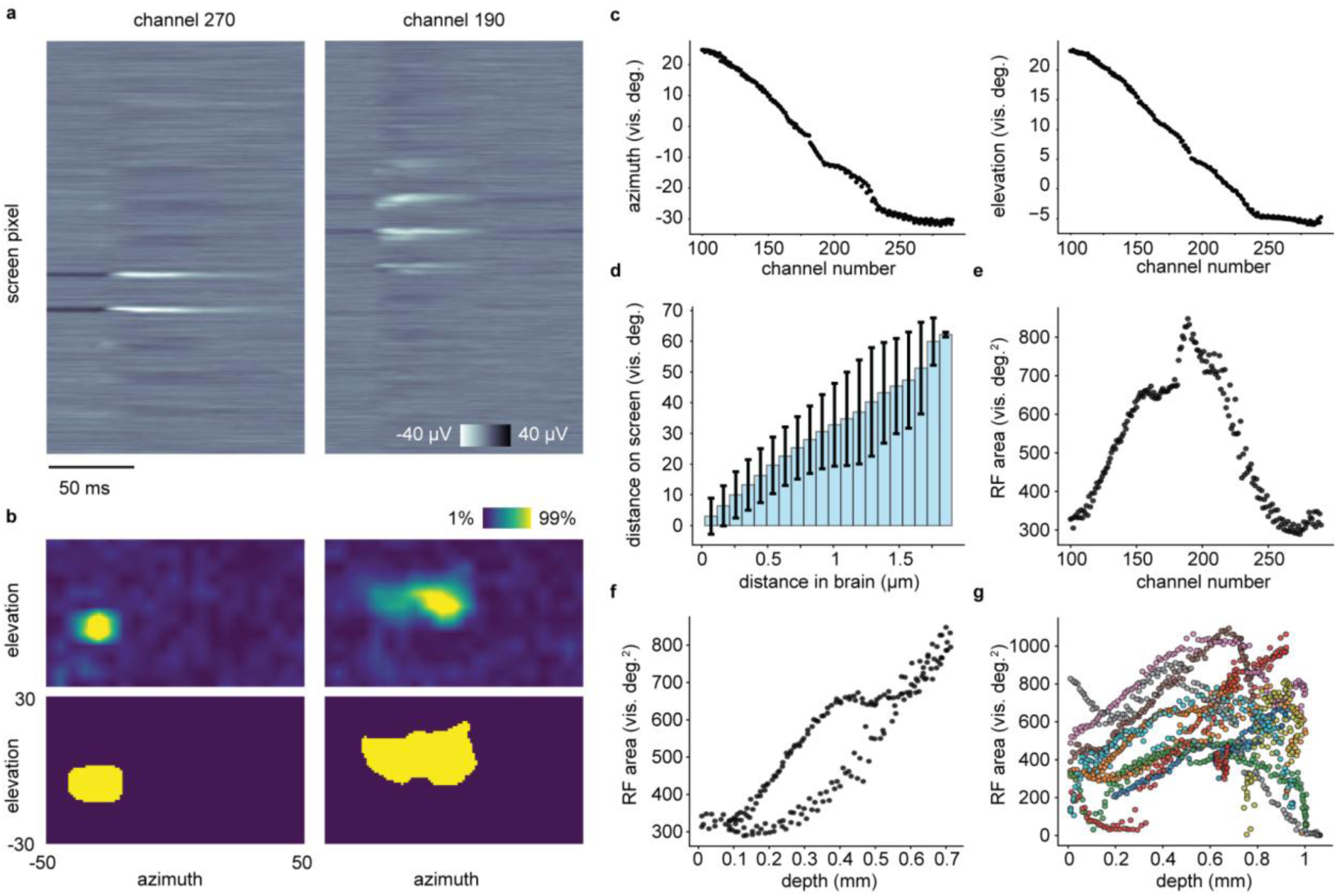
| Local field potential retinotopy and receptive fields. **a-b,** Two channels from probe shown in Fig. 4b. Left: more superficial channel, Right: deeper channel. **a,** Event triggered average LFP waveforms over time, triggered on a 5×5 visual degree sparse noise pixel turning from grey to white. **b,** Top: LFP event triggered average over screen position. Bottom: Estimated LFP receptive field (RF). **c**, LFP RF position on the screen in azimuth (left) and elevation (right), as a function of probe channel number, for probe in Fig. 4b. **d**, Pairwise distance of LFP channels in the brain and their RF positions on the screen. (n = 93945 channel pairs, 9 animals, Pearson’s R = 0.76, p = 1.0 × 10^-4^ from 10,000x bootstrap resampling). **e**, LFP RF area as a function of channel number for probe in Fig. 4b. **f**, LFP RF area as a function of depth in the medulla, for probe in Fig. 4b. **g**, LFP RF area as a function of depth in the medulla over experiments, showing an increase in area with depth (comparing data from 0-0.5 vs. 0.5-1 mm depths in the medulla, Mann-Whitney U: 155972, p = 8.51e-18, n = 1333 electrodes, 9 animals).

**Extended Data Figure 14.**
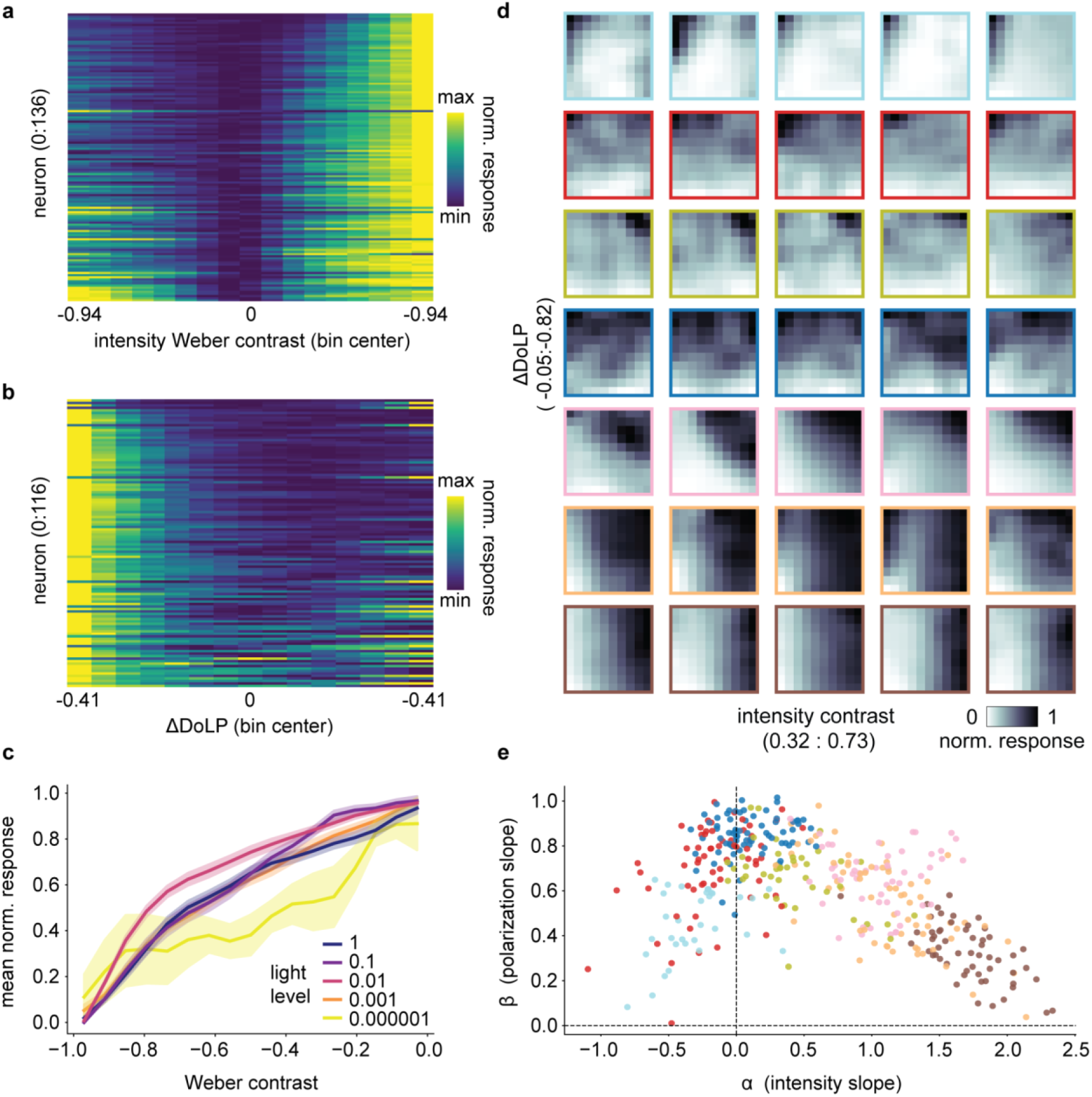
| Encoding intensity and polarization contrasts. **a,** Individual neuron response strengths as a function of intensity contrast (Methods). Neurons sorted by summed normalized response. **b,** As a, but for DoLP contrast. **c,** Response strength, averaged over neurons, for different contrasts and light levels. Background was set to black, and bars were delivered from black to 50% gray. (Light level 1 (full) to 0.000001 dropped by neutral density filters: n = 60,12,58,63,5 neurons, 3,1,2,3,1 animals per condition, data is mean ± s.e.m.). **d,** Example neural responses to int+DoLP stimulation (Fig. 5e-h). Outline colours match cluster identity in Fig. 5f. **e,** Distribution of additive model coefficients, for neurons in Fig. 5f-h. Colours match cluster identity in Fig. 5f.

**Extended Data Figure 15.**
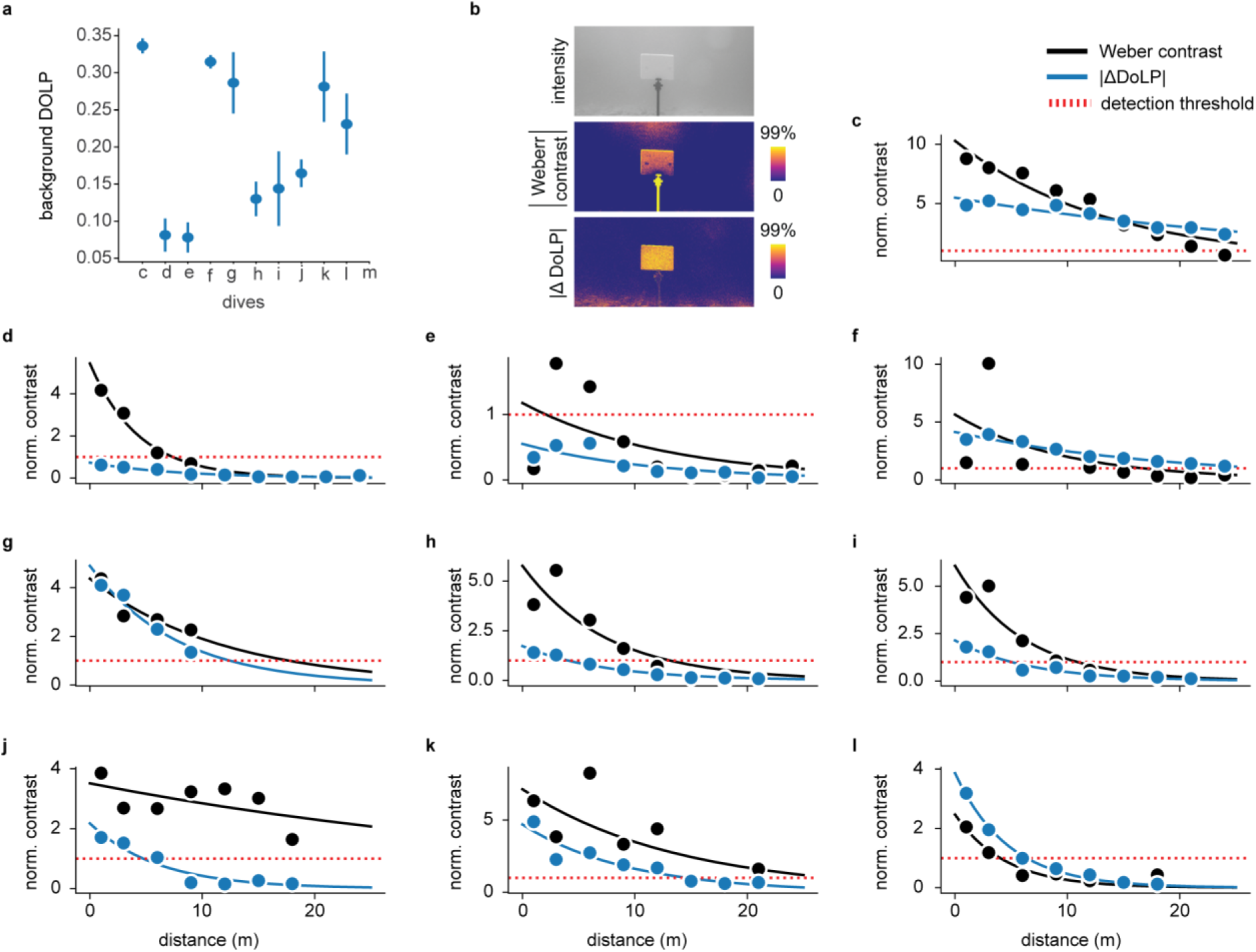
| Intensity + polarization contrast in the lab and underwater. **a**, DoLP levels recorded from the water column (background) across all dives. Vertical bars indicate ± 1 SD of background pixels. **b**, Intensity, absolute Weber contrast, and absolute ΔDoLP images of a white card, viewed facing the sun. For the bottom two images, colour scales are set from 0 to a value of 99% of data distribution (middle: 0::0.23, bottom: 0::0.14). Values below physiological detection thresholds were set to 0. **c-l**, Each plot shows Weber intensity and absolute ΔDoLP values as a function of distance for one dive (depth = 10 m). Values are normalized (divided) by the physiological detection threshold. Grey dotted line: physiological detection threshold (following normalization = 1). Estimated sighting distances (intensity/DoLP, in meters) c : 25(max)/ 25(max), d: 7.4/0, e: 1.7/0, f: 16.6/25(max), g: 17.7/12.4, h: 13.1/4.0, i: 10.9/5.0, j: 25(max)/4.7, k: 25(max)/14.6, l: 4.0/6.4

## Supplementary Video Description

**Supplementary Video 1: Two-photon Ca^2+^ imaging from the awake squid optic lobe.** Imaging plane in the OGL, neurons imaged during sparse noise visual stimulation. Video is motion corrected and denoised (Methods).

**Supplementary Video 2: Bouts of IGL Ca^2+^ wave activity are accompanied by atypical behaviour**

Left: Ca2+ imaging from the IGL (scale bar: 50 μm; playback is sped up 4×). Waves begin at approximately 50 s. The video is motion-corrected and denoised. Right: Time-aligned behaviour of the squid, synchronized with the calcium movie.

**Supplementary Video 3: 3D light sheet scan of the squid brain**

The squid brain was cleared and stained with TO-PRO-3 (nuclear marker, purple) and an anti-alpha-tubulin antibody (green; see Methods). The 3D image was acquired with voxel size of (X, Y, Z) = (0.65, 0.65, 2.5) μm and rendered using Imaris.

**Supplementary Video 4: Underwater video of squid in their natural habitat**

Video clips filmed while SCUBA diving in Seragaki bay, Okinawa, Japan. Depths = 1-5m. Squids are *Sepioteuthis lessoniana*. Note the caustic light fluctuations complicating intensity-based vision.

## Methods

### Experimental Animals

#### Tank systems setup & water quality

All research and animal care procedures were conducted in accordance with national laws and institutional guidelines and were approved by the OIST Animal Care and Use Committee (approval numbers 2019-244-6 and 2022-364). Juvenile squid (*Sepioteuthis lessoniana*, mantle length approximately 1.2-2.5 cm) of both sexes were raised from eggs in tanks connected to closed 1500 L seawater recirculation systems at the OIST Marine Science Station. Each circulation system possessed mechanical filtration, a protein skimmer, UV-filtration, and an MBBR (Moving Bed Biofilm Reactor) for biological denitrification (conversion of ammonia to nitrate). The systems had been previously N-cycled, so that the plastic carriers of the reactor could develop a biofilm allowing the concentration of NH_3_/NH_4_+, NO_2_, and NO_3_ to be kept well within optimal parameters (NH_3_/NH_4_+<0.1 ppm, NO_2_<0.05 ppm, NO_3_<50 ppm)^78^. Additionally, a 25% water change was performed biweekly using sand-filter natural seawater. The photoperiod was set to a 12-hour light/12-hour dark schedule, the temperature to 25°C and pH was kept within a range of 8.10-8.20. Salinity was maintained between 34.5 and 35.5 PSU, based on the salinity of the natural seawater collected at Seragaki, Okinawa, compensating evaporation by adding fresh water. To stabilise and maintain pH in the recirculation systems, alkalinity was brought to 9 dKH (3 dKH higher than the original natural seawater) by dosing NaHCO_3_ into the system. Calcium was kept at 425 ppm by dosing CaCl_2_ as needed^79^. Carbonates and calcium were always dosed on separate days to prevent precipitation. Salinity, temperature, and pH were monitored and adjusted daily, while alkalinity, calcium, NH_3_, NO_2_, and NO_3_, were measured weekly.

Both eggs laid inside open circulation breeding tanks at the OIST Marine Science Station and wild eggs collected from the field were used. After temperature acclimation, eggs were moved to floating baskets within dark square fiberglass tanks (115 cm x 55 cm x 45 cm, filled to 40 cm, water volume ∼250 L). Immediately after hatching, squid paralarvae were transferred from the incubator tanks system to the keeping tanks system, where they were group-housed with siblings in 11 L square glass tanks (19 cm x 29 cm x 20 cm, filled to 15 cm, water volume ∼8 L). The paralarvae were moved using transparent plastic containers, to minimise stress and avoid injuries from net use. Each keeping tank hosted a maximum of 25 hatchlings, approximately 3 individuals/L. Enrichment, such as artificial plants or coral rubble, was added to each tank, and a lid was placed over tanks with animals older than 1 month to prevent mortality due to jumping.

#### Squid husbandry

Immediately upon transfer to keeping tanks, squid were fed with live mysids (*Neomysis* spp.)^80^, sourced from mainland Japan and kept in large square tanks within a 1500 L recirculation system with brackish water (15-20 PSU). Mysids were provided to the squid *ad*-*libitum* for the first 7 days post-hatching (dph, approximately 5–7 mysids per squid per day), then twice daily from 8 to 15 dph (approximately 4 mysids per squid per day), and once daily from 15 to 30 dph (approximately 1–2 mysids per squid per day). Concurrently, the animals were trained to accept dead food, which was introduced on 1 dph and offered four times a day. As the squid grew, the defrosted dead food, consisting of a fish-shrimp mixed diet, was provided ad libitum, gradually replacing mysids as the primary food source. Live food was delivered until animals would accept 100% of defrosted food, typically by 30 dph. Squid were examined every morning for signs of stress (abnormal behaviour, excessive inking, protracted refusal of food, or incapability of hunting) and monitored for malnutrition, injuries or diseases throughout the day. Animals that appeared unhealthy were removed from the tanks and euthanized using ethanol. To perform euthanasia, the squid were anesthetized by submerging them in a mixture of seawater and ethanol at 1.5% (15 mL/L). Following anesthesia, more ethanol was gradually added until the concentration reached >3% (30 mL/L)^81^ and breathing ceased. Death was visually confirmed by observing the brain changing from transparent to white/opaque. Each keeping tank was syphoned three times a day to remove food remains and feces. The filters from the recirculation systems were cleaned every two weeks. All tools used for animal keeping were rinsed in fresh water after each use, in addition tweezers used for feeding were sterilised using ethanol after washing.

### Head Fixation

Prior to physiological experiments, animals were head fixed as follows: first, the squid was anesthetized using 1% ethanol in sea water. Once the squid was fully anesthetized (usually in 1 to 2 minutes), it was transferred to a surgery dish (a 15 cm diameter cell culture dish with a silicone floor) filled with oxygen-saturated natural sea water. A thin metal wire loop was used to briefly stabilize the animal on the silicone floor, and the skin above the OL was removed with forceps to expose cartilage. After briefly drying the cartilage with a Kimwipe, a drop of green silicone glue (WPI Kwik-Cast) was applied around the planned recording site (∼1.5 mm diameter). Subsequently, the rest of the exposed cartilage was covered with tissue adhesive (3M Vetbond) to reinforce the rigidity of the cartilage. Next, a stainless steel head plate was glued onto the reinforced cartilage using a UV-curing glue (3M Transbond XT Primer). The head plate had a 1.8 or 2.3 mm diameter circular hole through which electrode insertion or imaging was performed. Finally, the green silicone glue was carefully removed so that the glue-free recording site was exposed (Extended Data Fig. 1a). After head fixation, the squid was transferred to a recording tank (150×104×95 mm inner dimension; transparent acrylic) and fixed onto a 3D-printed elevated platform. For some recordings we gently placed the mantle in a transparent tube (5-8 mm inner diameter) to prevent large mantle movements without interfering with normal breathing (Extended Data Fig. 1c). Additionally, a plastic piece was positioned above the arms to prevent contact with the electrode or objective lens, while allowing free movement otherwise. For electrophysiological experiments, small holes (∼70 μm diameter) were made in the cartilage using a glass micropipette to allow Neuropixels probes to enter the brain. Special care was taken to avoid damaging the large vein on the OL surface when puncturing holes.

### Calcium indicator dye injection

After head fixation, a calcium indicator dye, Cal-520 AM (AAT Bioquest 21130), was introduced to the OL neurons using a pressure injection, using a similar method to previous work in octopuses^17^. First, artificial sea water (ASW) was prepared using the following recipe: 460 mM NaCl, 10 mM KCl, 40 mM MgCl2, 11 mM CaCl2, 10 mM Glucose, 10 mM HEPES, 2 mM glutamine. The injection solution contained 2.25 mM Cal-520, 10% DMSO, 2% Pluronic (Thermo Fisher P3000MP) and 0.05 mM Alexa Fluor 594 Hydrazide (Thermo Fisher A10438) in ASW. This injection solution was loaded into a glass micropipette connected to a pneumatic picopump (WPI PV820). The tip of the glass micropipette had 8-10 μm inner diameter and was beveled at ∼30° to puncture the cartilage without clogging. The solution was injected into the tissue by applying weak pulsatile pressure (∼2 psi for 1 second), every 30 seconds, for 5 to 10 times at the targeted depths. This procedure resulted in homogeneous labeling of the cells in ∼300 μm diameter. After the injection, a drop of transparent silicon glue (WPI Kwik-Sil) was applied to the cranial window, and a cover glass was secured onto the window using UV glue.

### Neural tracer injection

After head fixation, a lipophilic tracer dye, CM-DiI (Thermo Fisher C7000; 5 mg/mL in DMSO), was injected into the OL medulla via pressure injection. A glass pipette was inserted to a depth of 1000-1500 μm from the surface. One or two brief pressure pulses (∼2 psi for 500 ms) were applied to eject a small volume (up to a few nanoliters) of CM-DiI. Following the injection, the animal was released from head fixation and allowed to recover in a home tank for 1 day (sample #1) or 7 days (sample #2 and #3). The animal was then euthanized, and the brain was dissected for tissue clearing. 3D images acquired by light-sheet microscopy were visualised using Imaris (version 9.7, Bitplane) and analysed by a custom Python script.

### Visual stimuli

A custom-made system was used to control light intensity and polarization delivered to the animal (Extended Data Fig. 3a)^3,82^. We used a gamma-calibrated projector (Texas Instruments DLP3010EVM-LC), to backproject images onto a modified vertical alignment type liquid crystal display (VA-LCD) (EIZO S1504). The modification included decasing of the panel from the enclosure, mounting the panel in a custom frame, removing the front polarizer, and attaching a diffuser sheet (LEE Filters 250 Half White Diffusion) on the backside of the panel. With this modified VA-LCD, we could effectively change the degree of linear polarization (DoLP) at each pixel from near 0 to 1. The calibration result of EIZO S1504 is shown in Extended Data Fig. 3b. To quantify the angle dependence of intensity changes using our DoLP monitor, we captured images of the LCD screen, with the projector intensity set to maximum, using an intensity camera (Basler ace acA4112-30uc). Mean intensity values were calculated from pixel ROIs from different viewing angles relative to the camera. Changes in intensity were calculated by comparing the difference in intensity between fully polarized and fully unpolarized LCD settings, from regions of interest at viewing angles encountered by the squid in our experiments. This was performed with and without a seawater filled acrylic tank placed in between the LCD and camera (Extended Data Fig. 3c). We repeated this procedure with a polarimetric camera (Basler acA2440-75umPOL, with Sony IMX250MZR CMOS sensor) to test the dependence of measured DoLP changes on viewing angle (Extended Data Fig. 3d,e).

Visual stimuli were generated and rendered through custom Python scripts written with the Psychopy library^83^. Both projector and LCD operated at 60 Hz refresh rate. For calcium imaging experiments, the projected image size was 19.2×10.8 cm, and the squid was positioned 7.0 cm away from the screen, covering ∼108 degree horizontal visual angle. At the maximum intensity, the light level measured at the squid eye position was 0.08 W/m^2^. Stimuli included the following:

- **Linear polarizer rotation**: A linear polarizer film (Edmund Optics XP42HE-40) was mounted on a motorized rotation stage (Thorlabs ELL18) using a custom 3D-printed housing. The polarizer was rotated through 16 angles from 0° to 168.75° in 11.25° increments, presented in sequential order. At each angle, we delivered a full-field intensity stimulus using a projector, consisting of 4 seconds OFF followed by 4 seconds ON, before advancing to the next angle. After completing all 16 angles, the entire sequence was repeated 10 times.
- **Locally sparse noise**: white squares (100% intensity) presented on a black background (0% intensity), refreshed at 1 Hz. The square size was 2°, with center-to-center spacing of at least 9°. Each pixel on the screen was sampled at least 18 times.
- **Drifting gratings**: binary bars patterns with either (black 15° / white 5°) or (black 7.5° / white 2.5°) widths, presented at random orientations from 0° to 315° in 45° increments, drifting at 3 cycles per second. Each grating was displayed for 2.5 seconds without interleaving frames. Three intensity/polarization phases were tested:

- (1) intensity gratings on an unpolarized screen (DoLP = 0.003)
- (2) intensity gratings on a polarized screen (DoLP = 0.950)
- (3) DoLP gratings with constant intensity, in which bars with DoLP=0.003 were presented on a background with DoLP = 0.950. Each full trial (phases 1-3) was repeated 18 times.
- **Full-field stimuli**: Each trial included three stimulus phases:

- (1) DoLP =0.003 (unpolarized background), stepwise intensity changes (OFF for 4 seconds, ON for 4 seconds, OFF for 4 seconds, gray for 2 seconds) followed by a sinusoidal chirp (0.5 to 10 Hz frequency sweep in 8 seconds).
- (2) DoLP =0.950 (vertically polarized background unless otherwise noted), same intensity protocol as phase 1.
- (3) Constant intensity (100%) with stepwise changes in DoLP followed by a chirp. Each full trial (phases 1-3) was repeated 10 to 15 times. In a subset of experiments, the screen was rotated 90 degrees between consecutive recordings to sample both horizontal and vertical angles of polarization (Figure 2a). In another set of experiments (Extended Data Fig. 6h-j), the screen was positioned at three angles (22.5°, 67.5°, and 112.5°) to sample horizontal, diagonal, and vertical angles, respectively (Extended Data Fig. 6h). These angles were chosen to closely align with observed peak angle sensitivities (Extended Data Fig. 6f,g), and 45° offset from these. For initial electrophysiology experiments, the projected image was 19.0×10.7 cm, and the squid was positioned 7.0 cm away from the screen, covering a ∼105° horizontal and ∼ 60° vertical visual angle. In later experiments, the projected image was 24.0×13.5 cm, and the squid was positioned 6.3 cm away from the screen, covering a ∼125 horizontal and ∼70 vertical visual angle. Light levels measured at the squid eye position ranged from 0.0012W/m^2^ to 0.209W/m^2^. Stimuli included the following:
- **Locally sparse noise:** white and black squares (100% and 0% intensity, respectively) presented on a gray background (50% intensity), refreshed at 10 Hz. For DoLP polarization experiments, squares were presented with high DoLP (95% DoLP) and low DoLP (3% DoLP) squares presented on a 50% DoLP background. In both cases, squares spanned 5 visual degrees in azimuth and elevation. No two squares were allowed within 20 visual degrees of each other at the same time. Intensity and polarization trials were interleaved (blocks of 200 trials) to control for slow shifts in spike rate. In a subset of experiments, the screen was rotated 90 degrees between consecutive recordings to sample both horizontal and vertical angles of polarization.
- **Sinusoidal drifting gratings:** For direction selectivity experiments, sinusoidal gratings (black to white) were presented with random travelling directions (0° to 315° in 45° increments), spatial frequencies (0.024, 0.06, 0.18, 0.26, 0.84, 1.2 cycles/degree) and temporal frequencies (0, 1.5, 3, 7 Hz). Each grating stimulus was presented for 1.0 seconds.
- **Binary drifting bars**: For contrast sensitivity experiments, binary bars travelling downwards (foreground) were presented with spatial frequency 0.06 cycles/deg and temporal frequency 1.0 Hz, with randomized intensities (0% to 100%) or DoLP (9% to 95%) on a 50% intensity or DoLP background, respectively. Stimuli were presented for 0.5 seconds, followed by a 0.5 seconds blank frame (as background).
- **Binary singular bars**: For the combinatory luminance/polarization experiment, we presented trials of singular bars (full-screen width, 22 visual degrees height) moving downwards simultaneously in both the projector and LCD display (foreground), which were aligned in location and size (+-2 pixels). Background levels were set at 40% intensity for the projector and 95% DoLP for the LCD, with bars having random values (40% to 80% for projector, 94% to 9% DoLP). Trials were presented at 1s intervals.
- **Optic flow:** To confirm direction selectivity results, an optic flow stimulus was presented to the animal with dots of different diameters (0.48, 0.97, 1.94, 2.91 visual degrees) traveling across the screen in random directions (0° to 315° in 45° increments), speeds (5, 17, 34, 50 visual degrees/second) and contrasts (black/white dots vs mean intensity backgrounds). Single values of stimulus parameters were used per trial. Dot movements were confined to the XY plane, reproducing drifting grating directions. Each optic flow stimulus was presented for 1.0 seconds.
- **Locally drifting Gabors:** To confirm direction selectivity results, Gabor patches of fixed spatial frequency (0.77 cycles/deg) and temporal frequency (3 Hz) drifting at random directions (0° to 270° in 90° increments) were presented on a mean intensity background. The patches spanned 5 visual degrees in azimuth and elevation. No two patches were allowed within 20 visual degrees of each other at the same time. Each presentation lasted 0.25 seconds.

For binary drifting grating and singular bar experiments, we converted intensity values into Weber contrast as

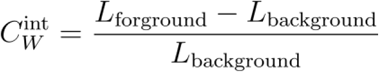

### Calcium imaging experiments

Calcium imaging was performed using a custom-built laser scanning two-photon microscope (based on Thorlabs DIY multiphoton microscope kit MM201). A 920 nm ultrafast laser (Spark Lasers ALCOR 920-2-XSight) provided the excitation beam and the emitted green fluorescence was detected by a GaAsP PMT (Thorlabs PMT2100) after passing through a dichroic mirror and a fluorescence filter (Semrock FF665-Di02 and FF03-525/50). The images were acquired using a 25X Objective lens (Olympus XLPLN25XSVMP2) with 512×512 pixels (0.52 μm per pixel) at 15.4 Hz unless otherwise specified. In a subset of experiments where we recorded spontaneous activity, we performed fast multi-z imaging using piezo objective scanner (Thorlabs PFM450E), which acquired 5 planes separated by ∼40 μm at 3.75 Hz. The total laser power after passing the objective lens was typically between 35 to 70 mW. The squid’s tank was integrated with a temperature control unit and a perfusion system connected to a large reservoir of oxygen-saturated natural sea water. To monitor the behaviour of the squid, it was illuminated by an infrared (IR) LED (Thorlabs M940L3) and filmed with a CMOS camera (Basler a2A1920-160ucPRO with an IR cut filter detached from the sensor and an IR pass filter attached to the lens) at 120 fps (Extended Data Figure 1b). In the same way, we also monitored the ipsilateral eye that received the visual stimuli with a second camera using a reflection by a hot mirror (Thorlabs FM01R). We employed a temporal multiplexing approach^84^ to present visual stimuli while performing two-photon imaging without crosstalk. The LED in the projector was connected to an external current driver (OpenEphys Cyclops^85^) and was pulsed in synchrony with the two-photon laser scanning, so that the projector illumination was active only during the fly-back period of the resonant scanner operating at 8 kHz. Only the blue LED (peak intensity at around 480 nm) was used throughout the calcium imaging experiments reported here. The synchronization between imaging, behavioural filming and visual stimulation was handled by a DAQ device (National Instruments PCIe-6353) and a custom LabVIEW script. To accurately determine the depth of recorded cells within the OL cortex, we acquired a z-stack image around the recording site (typical imaging volume: (X, Y, Z) = (532, 532, 250) μm). The template image for each recording (averaged over all frames) was aligned to the z-stack using a template matching algorithm implemented in OpenCV. In the z-stack image, the boundary between the PXL and IGL was manually annotated and used as a reference to determine the depth of each cell in the recorded movie. This approach allowed accurate depth determination along the curved surface of the OL.

### Electrophysiological experiments

Neuropixels recordings (version 1.0) were performed using SpikeGLX software (version 3.0), sampling at 2.5 kHz (for LFP signals) and 30 kHz (for extracellular spike signals). Squid were filmed simultaneously from a side view contralateral to visual stimuli at 150 fps. For a subset of experiments, the eye ipsilateral to the visual stimulus was filmed using a hot mirror. Videos were synchronised to electrophysiological recordings by sending a 150 Hz transistor-transistor-logic (TTL) signal from an Arduino to trigger camera frame exposure and to log TTL time using SpikeGLX. Visual stimuli were synchronised by presenting a small flickering square (black/white for projector stimuli, high DoLP/low DoLP for LCD stimuli) in a corner of the projected image. This region was covered so as to not be visible to the squid. The flickering timings were captured by amplified photodetectors (Thorlabs, PDA100A2), and logged using SpikeGLX. During experiments, fresh oxygen-saturated seawater was circulated through the recording tank.

### Acute slice preparation

Cross-sectional slices of the optic lobe were obtained from squid aged 25–45 days post-hatching (dph). Animals were anesthetized using 4% ethanol and killed by decapitation. Both optic lobes were quickly dissected and affixed to the stage of a vibrating microtome (Leica VT1000S or Dosaka NeoLinear Slicer Model MT) using superglue, with a 2% agarose block supporting the posterior side. Slicing was performed in a cold (4°C) bath of low-calcium artificial seawater (“cutting” ASW; in mM: NaCl 402.5, KCl 10, CaCl₂ 0.5, MgCl₂ 53, MgSO₄ 2, KH₂PO₄ 0.5, NaHCO₃ 30, glucose 10, HEPES 10; pH 7.35; 975 mosmol/kg), continuously bubbled with 95% O₂ / 5% CO₂^86^. Transverse slices were cut 400 μm thick and transferred to a holding chamber containing cold (4°C) “recording” ASW (same composition as above, but with 11 mM CaCl₂ and 43 mM MgCl₂, and 12 mM MgSO₄) for ∼60 minutes prior to recording. During experiments, slices were submerged and continuously perfused with recording ASW at 1–3 ml/min at room temperature. To stabilize the tissue, slices were secured in place using a U-shaped “harp.”

### Whole-cell patch clamp & two-photon microscopy in brain slices

Optic lobe cells were loaded with fluorescent dye via the whole-cell patch clamp method^87^. Patch pipettes (Borosilicate glass, O.D.: 1.5mm, I.D.: 0.86mm, Sutter Instruments) were pulled using a horizontal puller (Sutter P-97; tip resistance 1.5-3 MΩ). Patch pipettes were back-filled with an internal solution containing (in mM): 500 K-gluconate or CsCl, 10 NaCl, 4 MgCl₂, 3 EGTA, 20 HEPES, 2 Mg-ATP, 0.2 Na-GTP, and 0.05-0.2 Alexa Fluor 488 or 594 (Thermo Fisher Scientific). The solution was adjusted to pH 7.4, 900 mosmol/kg^86^. After achieving whole-cell configuration, the dye was allowed to diffuse into the cell for at least 10 min and up to 30 min before image acquisition. The patch pipette was left in place throughout acquisition. In some cases, the extracellular space was visualised via local perfusion of 0.1 mM AF488 (in recording ASW) through a separate glass electrode.

Images were acquired live on a Nikon AX RMP two-photon microscope using a NIR Apo 60x NA 1.0 water immersion objective (Nikon) at 920 nm (AF488) or 850 nm (AF594) excitation wavelength. Multiple volume scans covering the whole cell were taken at 1024 x 1024 pixels, 1 μm axial steps, 4x zoom, 4-8x line average, resulting in a voxel size of 0.0719 x 0.0719 x 1 μm (XYZ). Additionally, to determine the location of the cells within the acute slice, bright field images were taken on a digital CMOS camera (Hamamatsu, ORCA-spark, C11440-36U) at 60x and 10x magnification (Plan Fluor 10x NA 0.8W objective, Nikon).

In some experiments, 1 mM Glutamate dissolved in HEPES-buffered AWS (same as “recording” ASW, but with 0.5 mM NaHCO₃, and 25 µM Alexa Fluor 594 for visualization) was focally applied through a glass pipette (∼ 1 µm diameter) using a Picospritzer at 6-8 psi for 20-50 ms. The spread was monitored by simultaneously recording the Alexa Fluor 594 dye and minimized by adjusting the duration of the puff. Neural responses were recorded in voltage clamp mode while holding the cell at −55 mV.

### Magnetic Resonance Imaging

For the MRI scan in Fig. 1b, a 52 DPH squid was euthanized using ∼3% ethanol, then fixed in 4% paraformaldehyde (PFA) prepared with seawater. The fixed squid was washed with PBS and immersed in 3 mM Prohance in PBS for 2 days at 4C, then transferred to Fomblin (YL VAC 06/6, Solvay) for scanning. MRI scans were performed on a horizontal 11.7 T Biospec, using a 35 mm inner diameter quadrature volume RF coil (Bruker, Ettlingen, Germany). The sample was scanned using a 3-D FLASH sequence, with a 25 µm isotropic spatial resolution. We used the following acquisition parameters: repetition time= 60 ms, echo time = 11.3 msec, Flip Angle = 50, field of view = 15×14×10 mm, matrix size = 600×560×400, NA = 6. Total acquisition was 22h, 24min.

### Transmission electron microscopy

Following euthanasia, eyes were removed and fixed with primary fixative (2.5% glutaraldehyde and formaldehyde in 0.1 M sodium cacodylate buffer (pH 7.4)). Fixed samples were washed in 0.1 M cacodylate buffer and postfixed with 1% OsO4 + 1.5% KFeCN6 for 1 hr. Following postfixation, samples were then washed in water (2 x 10 min), 50 mM Maleate buffer (MB) (1 x 10 min, pH 5.15), and incubated in 1% uranyl acetate in MB for 1 hr. Next, samples were washed in MB (1 x 10 min), followed by water (2 x 10 min), and subsequently dehydrated in grades of alcohol (10 min each; 50%, 70%, 90%, 2 x 10 min 100%). Following dehydration, samples were put in propyleneoxide for 1 hr and infiltrated overnight in a 1:1 mixture of propyleneoxide and TAAB Epon (TAAB Laboratories Equipment Ltd). The following day, samples were embedded in TAAB Epon and polymerized at 60 °C for 48 hrs. Ultrathin sections (50-80 nm) were then cut on a microtome (Reichert Ultracut-S), picked up onto copper grids, stained with lead citrate, examined in a JEOL 1200EX Transmission electron microscope, and images were recorded with an AMT 2k CCD camera.

### Calcium imaging data processing

The recorded calcium imaging movie was processed using a custom Python pipeline for motion correction and denoising. First, rigid motion correction was performed using the NoRMCorre algorithm implemented in the CaImAn library^88^. The rigid-corrected movie was then denoised using DeepInterpolation, a deep neural network-based algorithm^89^. We used the pre-trained “Two-photon Ai93 excitatory line DeepInterpolation network” provided by the authors. The output was subsequently subjected to non-rigid motion correction using NoRMCorre, and the computed transformations were applied to the original rigid-corrected movie. This non-rigid corrected movie was further denoised using DeepInterpolation. Denoised movies were used solely for visualisation and automated cell detection, while fluorescence signals were computed from the original non-rigid corrected movie, to avoid potential artifacts introduced by denoising. Large amplitude brain movements were detected by computing the cross-correlation value between the template (mean over all frames) and each frame. Frames with low correlations were excluded from further analysis.

To detect candidate ROIs corresponding to individual cells, we used Suite2p^90^. We then manually curated the predicted ROIs using ImageJ^91^ and its ROI Manager tool. In cases where multiple recordings were made from the same imaging area, ROIs identified in different recordings were mapped together. To achieve this, we prepared template images from each recording and used SIFT feature detection along with the FLANN matching implemented in OpenCV to align them. These automatically mapped ROIs were manually validated and corrected.

The fluorescence signal for each cell was calculated by averaging the pixel values within its ROI and correcting for neuropil contamination^92^. The corrected somatic fluorescence signal was estimated as:

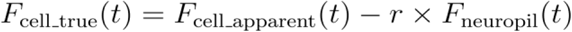

where r = 0.5 in our analysis. The neuropil signal, *F_neuropil_*(*t*), was obtained by averaging pixel values within 15 µm distance surrounding the cell outline, excluding all identified cell ROIs.

### Electrophysiology data processing

In SpikeGLX, a separate recording was acquired per visual stimulus used in an experiment. The AP bands (300 Hz High pass filtered data, 30kHz sampling frequency) of individual recordings were temporally demultiplexed, filtered for transient electrical artifacts (such as those caused by jerking movements of the animal), and concatenated into a continuous recording using the CatGT tool (https://github.com/billkarsh/CatGT). To identify spiking events, and assign them to individual units, the concatenated datasets were spike-sorted using the Kilosort library (Kilosort 4.0)^93^. The output clusters of the automated sorting were manually curated using the phy library (https://github.com/cortex-lab/phy), on the basis of waveform shape, autocorrelogram, cross correlogram, and interspike interval violations. Kilosort often assigned spikes to the residual of a waveform. Putative double-counted spikes were removed from the curated units when two spiking events were detected within a 0.16ms interval.

### Eye tracking

To track eye movements post-experimentation, we trained separate single-animal SLEAP^94^ models for each camera setup: electrophysiology ipsilateral eye, electrophysiology contralateral eye, calcium imaging ipsilateral eye, and calcium imaging contralateral eye. The model skeleton was designed to capture the boundary of the lens, eyelid, and eye muscle. For each model, 20 frames were manually labeled from each video used in both training and subsequent analysis. Skeleton centroids, matching the center of the squid lens, were used to quantify the location and movement of the squid eye.

#### Analysis

Probability density distributions were computed by dividing the number of frames in which a centroid occupied a given pixel by the total number of frames. Images were min-max normalized for display. Eye drift was quantified as the 2D Euclidean displacement of the centroid over time relative to its mean position, and converted from pixels to millimeters using a size calibration factor.

### Tissue clearing and light sheet imaging

Following Neuropixels recordings or neural tracer dye injections, the animal was euthanized by gradually increasing ethanol concentration to 5% in sea water. The brain was dissected and fixed in 4% paraformaldehyde in PBS at 4°C overnight. Dissected brain tissue was cleared using a modified BABB method^95^. First, the tissue was dehydrated in gradient concentrations of methanol (30%, 50%, 80%, 100%; each for 30 to 60 minutes at RT). The solution was refreshed with 100% methanol and incubated overnight at RT. On the next day, the sample was rehydrated in gradient concentration of methanol (80%, 50%, 30%) and washed with a wash buffer (5X SSC, 20 mM citric acid, 0.01% Tween-20 in water) twice. The tissue was then incubated in TO-PRO-3 nuclear staining dye (Thermo Fisher T3605; 2 μM dissolved in wash buffer) for 12-24 hours at 37°C. On the next day, after washing three times with a wash buffer, the tissue was embedded in a 1% agarose gel (nacalai tesque 01163-76) using a custom-made mold. Embedding in the agarose prevented sample movement during imaging. The tissue was dehydrated in gradient concentrations of methanol (30%, 50%, 80%, 100%), followed by overnight incubation in 100% methanol. On the last day, the sample was incubated for 6 hours in 1/3 methanol and 2/3 dichloromethane solution, then in 100% dichloromethane. The sample was transferred to the BABB solution and was cleared completely. We imaged samples using a custom-built light-sheet microscope, as described previously^31^. TO-PRO-3 was imaged with a 640 nm excitation laser and 692/40 nm bandpass filter, and CM-DiI was imaged with a 532 nm excitation laser and 593/40 nm bandpass filter.

For whole-brain immunostaining (Supplementary video 3), we followed the iDISCO method^96^ with the following conditions. Primary antibody: alpha-tubulin antibody (Sigma T6793) at 1/50 dilution, incubation at 37°C for 72 hours. Secondary antibody: Mouse IgG2b with Alexa Fluor 594 (Jackson laboratory 115-587-187) at 1/40 dilution, incubation at 37°C for 72 hours.

### Brain registration and probe localization

To construct a reference atlas of the *S. lessoniana* OL, we cleared, stained, and imaged a squid brain following the procedure described above. We then cropped the OL and neighboring central brain, downsampling to (X, Y, Z) = (10, 10, 10) μm. We used 3D Slicer^97^ to manually annotate 6 major regions in this 3D image: outer granular layer (OGL), plexiform layer (PXL), inner granular layer (IGL), medulla, peduncle lobe (PDL) and the central brain (CB). 3D images of individual squid brains were mapped to this reference brain, utilising the ANTs library^98^. Neuropixels probes were localised as described previously for octopuses^31^. The number of channels located in the brain for each probe was identified by computing a cross-correlation matrix between LFP signal channels. Channels outside the brain (in sea water) showed strong correlations with each other but weak correlations with channels inside the brain (Extended Data Fig. 10c). Brain registration quality was assessed by comparing the warped brains against the reference atlas for structural deformities (Extended Data Fig. 11e). Probe registration quality was assessed based on the directionality and continuity of probe channel (insertion site to probe tip), as well as the inter-channel distance distribution following registration (Extended Data Fig. 11f).

### Morphological reconstruction of neurons

Following image acquisition in whole-cell patch clamp experiments, Individual volume scans were manually stitched in ImageJ^99^ and combined via linear blending. The composite image was denoised with a 2D median filter (radius = 1 px) and gamma transformed (gamma = 0.25-0.5). Cell morphologies were semi-automatically reconstructed using the SNT toolbox in ImageJ^100^ and exported as SWC files for further analysis. Traces were further aligned to the major axis of the local cell island which is oriented perpendicular to the optic lobe surface. To visualise cell morphology, a mask around the tracing was created using the “Fill” function in SNT and applied to the original image. OGL and IGL neuron traces were further normalized to the thickness of the plexiform layer, which scales with the age and size of the animal. The thickness of the plexiform layer is strongly correlated with that of the IGL (Pearson r = 0.68, p < 0.001). Branch-points were detected as vertices with degree > 2. We found that regions with high branch-point density corresponded to areas of putative pre- and postsynaptic endings, both from visual inspection of the underlying morphology exhibiting varicose and filopodia-like structures, and in the case of postsynaptic endings, responsiveness to focally applied glutamate (data not shown), though further ultrastructural confirmation is required. For OGL neurons, a distinct peak of high-density branch-points close to the OGL boundary overlapped with the nerve endings of photoreceptors, which appeared during shadow imaging as filled, cylindrical compartments, and were assumed to be postsynaptic terminals. For IGL neurons, branching occurred both within the plexiform layer and in the Medulla. Given the supposed connectivity between IGL and OGL neurons, we assumed any cluster of branch-points within the plexiform layer to be putatively postsynaptic.

The distance from putative postsynaptic terminals to the soma was measured along the neuron processes from the branch-point to the centre of the soma. To estimate the radius of neurites, cross-sections along the primary neurite originating from the soma were taken and modelled as a boxcar function convolved with the point spread function (PSF) of the imaging system (measured empirically using 0.1 μm fluorescent beads, ThermoFisher):

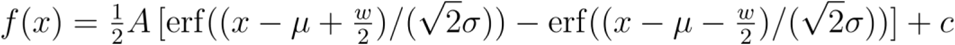

Where *A* = amplitude, μ = centre point, w = width, σ = standard deviation of PSF, and c = offset. The lower bound for the width was set to the theoretical diffraction limit of 0.31 μm.

The length constant of an ideal, cylindrical compartment is given by

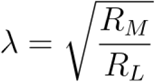

Where the membrane resistance R_M_ is given by

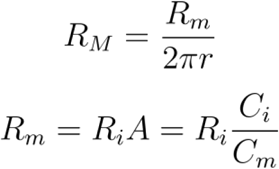

Where *R_m_* is the specific membrane resistance, r the radius, R_i_ the input resistance, A the area of the compartment, *C_i_* the input capacitance, and *C_m_* = 1 µF/cm^2^, the specific membrane capacitance^77^. The longitudinal resistance *R_L_* was calculated as

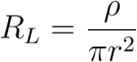

Where *ρ*=30 Ωcm is the axoplasmic resistivity^75,76^. The input resistance was calculated from the steady-state current, *I_ss_*, of voltage steps of −20 to +20 mV around resting membrane potential, which showed minimal voltage-dependent currents, accounting for the access resistance *R_a_*:

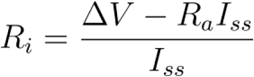

The input capacitance was similarly estimated as charge over voltage change:

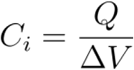

Where charge *Q* was calculated by integrating the current over the time span of the voltage change.

### Calcium imaging analysis

#### Receptive field (RF) analysis

Responses to locally sparse noise stimuli were recorded from both OGL and IGL. For each pixel location (i, j), the event-triggered average from each ROI was calculated as:

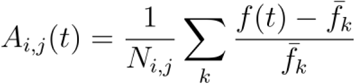

where f(t) is the neuropil-corrected fluorescence signal at time t, *N_i,j_* is the number of trials for location (i, j), *f̅_k_* is the baseline fluorescence averaged over a pre-stimulus window (−2 to 0 seconds) for trial k. The average response for each location (i,j) was quantified by integrating over a 1.0-second window following stimulus onset:

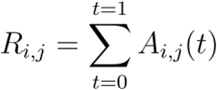

The resulting response map *R_i,j_* was z-scored across all pixels to identify significantly responsive regions. Pixels with z-scores greater than 3 or less than –3 were considered to show significant positive or negative responses, respectively. For each case, a binary mask was created by thresholding the z-scored map, and the connected component containing the peak was defined as the receptive field. The centroid and area of each receptive field were then computed.

#### Clustering of deep retina neurons

Responses to full-field visual stimuli were recorded from both OGL and IGL. The fluorescence signal from each ROI was first z-scored within each trial as:

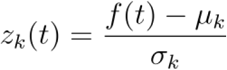

where f(t) is the neuropil-corrected fluorescence signal at time t aligned to the onset of the trial, *µ_k_* and *σ_k_* are the mean and standard deviation of the signal across the k-th trial. To assess response reliability, we computed the quality index (QI) for each neuron:

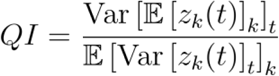

where 𝔼[]*_x_* and Var[]*_x_* denote the mean and variance across the indicated dimension x, respectively ^101^. Neurons with QI values greater than 0.25 were retained for downstream analysis. Additionally, neurons with large trial-to-trial variability were excluded (90th percentile of Var [*z_k_*(*t*)]*_k_* greater than 2.0).

After filtering, the mean z-score of each neuron across trials was computed as

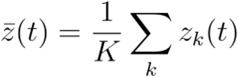

and was assembled into a response matrix X of shape N x T, where N is the number of neurons and T is the number of timepoints. Each row of X was mean-centered, followed by column-wise normalization using StandardScaler() from the scikit-learn library. Dimensionality reduction was performed using sparse principal component analysis (sPCA), also implemented in scikit-learn. sPCA was applied separately to each of the three stimulus phases, yielding 30 components in total. The resulting sparse component matrix S (shape: N x 30) was used for Leiden clustering to identify functional neuronal types. A neighborhood graph was constructed using pp.neighbors() function in the Scanpy library, followed by clustering via tt.leiden(). The algorithmically derived clusters were further curated manually by examining the mean response and the distribution of functional indices (described below) within each cluster. Clusters with overlapping properties were merged. Finally, we computed a distance matrix using cosine distance between cluster-mean responses and sorted the clusters using hierarchical clustering, using the single linkage method.

#### Functional indices to characterize clusters

To characterize the functional cell types identified above, we computed five indices from the mean z-scored calcium signal *z̅*(*t*). α₁, α₂, and α₃ are activation indices for full-field stimulus phases 1, 2, and 3, respectively. Each was computed by subtracting the mean fluorescence signal during the stimulus period (t = 0 to 4 seconds; light ON or low DoLP) from the pre-stimulus baseline (t = −2 to 0), followed by normalization to the range [−1, 1] while preserving sign. The high-frequency activation index, β, was computed by fitting a linear function to the calcium signal from t = 10 to 18 seconds (temporal chirp period) and extracting the slope, which was then normalized to [−1, 1] with sign preserved. The correlation index, ρ, was computed as the Pearson correlation between the responses in phases 1 and 2, using the cropped time window t = −2 to 8 seconds. These indices were computed for each neuron, and the cluster-wise means are shown in Fig. 1h.

#### Spatial distribution of OL cortex neuron types

To characterize the spatial distribution of functional cell types within individual recordings, we first selected recordings with more than 150 neurons that passed filtering criteria (n = 4 recordings). For each recording, we extracted cluster IDs that included more than 30 cells and computed Ripley’s L function, *L_r_*, (implemented in the pointpats library) based on the coordinates of cells belonging to each cluster. We then plotted *L_r_* – *r* against *r*, where *r* is the radial distance (Fig. 1k). We compared the resulting curves to the 2.5th–97.5th percentile range of simulated complete spatial randomness (CSR). Most curves fell within this range; in a few instances, parts of curves slightly exceeded the upper bound, suggesting spatial clustering. This may reflect a tilted imaging plane relative to the OL cortex which has a heterogeneous distribution of cell types along the depth axis.

As an alternative spatial statistic, we computed the k-th nearest-neighbour entropy using the same dataset. For each cell, entropy was calculated based on the cluster identities of its six nearest neighbours, and the mean entropy across all cells was computed. We compared this value to a null distribution generated by randomly shuffling cluster IDs while preserving cell positions. In all four recordings, the observed entropy was not significantly lower than the shuffled distribution (p > 0.05).

To assess the spatial distribution of functional cell types along the OL cortex depth axis, we selected recordings in which depth could be accurately determined (n = 792 cells from 5 recordings made in the OGL). For all cluster IDs containing more than 8 cells within this dataset (Cluster 5 and 14 were excluded by this criteria), we performed a Kruskal–Wallis H-test and found significant differences in depth distributions across clusters (H = 129.6, p = 2.06 × 10⁻²²). We then conducted post hoc pairwise comparisons between all cluster pairs using Dunn’s test, with Holm–Bonferroni correction for multiple comparisons (Fig. 1m).

#### Analysis of monitor-flip experiment

To investigate the polarization angle specificity of OL cortex neurons, we recorded calcium activity evoked by a full-field visual stimulus presented with two orthogonal screen orientations (see above; OGL: n = 548 neurons, 4 animals; IGL: n = 790 neurons, 3 animals). We focused on the time window from t = −2 to 8 seconds in phase 2 (intensity changes on a polarized background) and phase 3 (DoLP changes at constant intensity) to characterize the polarity of responses. Z-scored responses were computed as described above, and we filtered neurons that showed a significant response in ΔDoLP stimulus under either or both horizontal and vertical screen orientations. Specifically, we selected neurons with QI values greater than 0.25 and response amplitudes (defined as the difference between the 95th and 5th percentiles after applying a high-pass filter at 0.08 Hz) exceeding 0.6. This filtering yielded a dataset of DoLP-responsive neurons (OGL: *n* = 214; IGL: *n* = 297). Dimensionality reduction and Leiden community clustering, as described above, revealed 12 putative functional types differentiated by the polarity, amplitude, and kinetics of their responses (Fig. 2b, Extended Data Fig. 7a–c). To quantify polarization-angle specificity (or invariance), we defined the degree index (DI), calculated as the Pearson correlation between responses (t = –2 to 8) in phase 3 under vertical and horizontal screen orientations. A negative DI indicates anti-correlated activity, reflecting angle-specific tuning. A positive DI indicates tuning to DoLP changes that is invariant to polarization angle.

#### Analysis of linear polarizer rotation experiment

Responses to intensity stimuli at 16 polarization angles (0° to 168.75°; 11.25° increments) were recorded from the OGL and IGL (OGL: 5 recordings from 3 animals; IGL: 2 recordings from 2 animals). For each neuron, we computed the mean response across the full rotation sequence, and included neurons with a QI value greater than 0.2 in the analysis. Dimensionality reduction, Leiden community clustering, and manual curation were performed as described above (see “Clustering of deep retina neurons”), yielding 11 putative cell types defined by their polarization-angle tuning and response kinetics (Extended Data Fig. 6f,g).

#### Analysis of 45-degree monitor experiment

We recorded calcium activity evoked by a full-field visual stimulus presented at three screen orientations in consecutive sessions (session 1: horizontal, session 2: 45° offset, session 3: vertical; OGL: 3 recordings from 2 animals; IGL: 2 recordings from 2 animals; Extended Data Fig. 6h). Z-scored responses were computed as described above, and neurons were filtered based on the following criteria: (1) QI value greater than 0.2, and (2) response amplitude (defined as above) exceeding 0.9 in either (session 1, phase 3) or (session 3, phase 3). The latter criterion selected neurons that were sensitive to polarization angle or degree. This filtering yielded 264 neurons (OGL: n = 223; IGL: n = 41). To quantify the similarity between phase 1 and 2 (intensity change with unpolarized vs. polarized screens), we computed the cosine distance between the z-scored activity traces for the two phases and compared these distances across sessions (Extended Data Fig. 6i). To quantify the responsiveness to DoLP stimuli, we compared the response amplitudes (defined as above) for phase 3 across sessions (Extended Data Fig. 6j). For both analyses, we used the maximum value from session 1 or session 3, because neurons often responded preferentially in one of these two conditions.

#### Direction and orientation selectivity analysis

Responses to drifting grating stimuli were recorded from the OGL and IGL (OGL: 4 recordings from 3 animals; IGL: 7 recordings from 4 animals). For each angle θ, the event-triggered average from each ROI was computed as

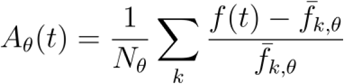

where f(t) is the neuropil-corrected fluorescence signal at time t, *f̅_k,θ_* is the baseline fluorescence averaged over a pre-stimulus window (−10 to 10 seconds) for trial k at angle θ. The average response for each angle was quantified by integrating over a 2.5-second window following stimulus onset:

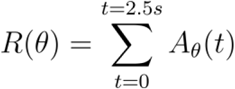

To quantify the direction selectivity, global direction selectivity index (gDSI) was computed:

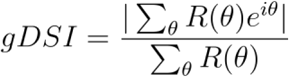

Likewise, to quantify the orientation selectivity, global orientation selectivity index (gOSI) was computed:

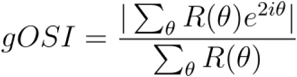

The statistical significance of direction selectivity was evaluated using a permutation test. Trial labels were randomly shuffled (n = 10,000 permutations) to construct a null distribution of gDSI values for each neuron. The p-value was computed as the fraction of shuffled indices that exceeded or equaled the observed value. A neuron was classified as significantly direction-selective if it met all of the following criteria: (1) p-value is less than 0.05 (2) *R*(*θ*) of the preferred direction was greater than 0.04 and (3) QI value is greater than 0.08. Orientation selectivity significance was assessed using the same criteria and procedure, substituting gOSI in place of gDSI.

#### Analysis of spontaneous activity

Spontaneous calcium activity in the OL cortex was imaged while the squid rested in complete darkness. Each recording lasted approximately 30 minutes. In a subset of experiments, simultaneous recordings from both the OGL and IGL were performed to compare activity between the two regions. In multi-plane imaging experiments, rigid motion correction was performed using a z-plane containing the OGL, and the same transformation was applied to the IGL. Putative ROIs were manually identified, and calcium signal traces from each ROI were resampled at 10 Hz and smoothed using a Gaussian filter (sigma = 3). Transient calcium spikes were detected using the find_peaks() function from the scipy library. The number of detected spikes was converted to spikes per minute and pooled across datasets.

Wave-like propagation of calcium signals was identified by manually reviewing the video recordings. To quantify bouts of repeated wave activity, ROIs were manually annotated, and the propagation speed and direction were analysed based on the timing of peak calcium activity within each wave.

### Electrophysiological analysis

#### Neural receptive field estimation

Receptive fields (RFs) of neurons recorded electrophysiologically were estimated using a locally sparse noise stimulus (see above). For each neuron, the event-triggered average (ETA) response and a null (trial shuffled) ETA were computed using a time window of 30-130 ms post stimulus. ETAs were computed separately for different stimulus types (white vs. black squares, or low vs. high DoLP squares). To extract RF size and shape, the ETAs were linearly upsampled by a factor of 5 (1 vis. degree/pixel) and smoothed using a Gaussian filter (sigma = 1 pixel). The null mean was subtracted from the response and the resulting matrix was thresholded (threshold: 5 SD of null). Estimated RFs were curated manually, with inclusion criteria being the presence of a true ETA peak, a good perceptual match between the RF estimate and the ETA, and for analysis of the RF size and location, a lack of RF overlap with the boundary of the visual display.

#### Receptive field distances on screen and in brain

For each neuron with a receptive field, the Euclidean distance between its receptive field location on the screen and that of other neurons with receptive fields recorded in the same session was computed. These distances were plotted against the corresponding Euclidean distances between neuron pairs in the brain, calculated from their positions after atlas registration.

#### Peri-stimulus time histograms

For neurons in Fig. 4e, peristimulus time histograms (PSTHs) were computed using trials in which the stimulus was active at the peak location of each neuron’s receptive field. Spike times were aligned to stimulus onset. Spike counts were accumulated in 10 ms non-overlapping bins and converted to instantaneous firing rates.

#### Visual modulation index (vmi)

To quantify the degree of activity modulation for each neuron with a visual receptive field, we computed a visual modulation index based on the responses to the locally sparse noise stimulus as:

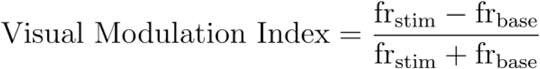

where fr_stim_ is the mean firing rate of the neuron at the peak location of its receptive field, computed from trials in which the stimulus was active at that location. Fr_base_ is the mean firing rate computed from trials in which no square within the receptive field was active, either in the current or the immediately preceding trial.

#### Waveform classification

Spike waveforms were extracted from extracellular recordings and aligned prior to shape classification. For each waveform, the sample corresponding to the global maximum or minimum (whichever had the larger absolute amplitude) was shifted to the centre of the waveform. Aligned waveforms were then classified based on the sequence of voltage extrema. Local maxima (peaks) and minima (troughs) were detected with a threshold set to 20% of the largest absolute extremum of the waveform. Based on the number and temporal order of detected extrema, waveforms were categorized into the following classes:

- positive: waveforms with a largest amplitude positive peak.
- negative monophasic: waveforms with a single prominent negative trough.
- negative biphasic: waveforms with one prominent peak and one prominent trough, further sub-classified as peak-before-trough or peak-after-trough based on their temporal order.
- negative triphasic: waveforms with three prominent extrema.

#### Direction selectivity analysis

For sinusoidal drifting gratings, neural direction selectivity was assessed as follows: For each neuron, the spike counts for each trial combination (spatial frequency, temporal frequency, and direction) were computed and summed across temporal frequencies and the four lowest spatial frequencies (there was no variation in firing for high spatial frequencies and were thus excluded from analysis), and were used to identify the peak direction. To quantify direction and orientation selectivity, a global direction selectivity index and global orientation selectivity index (as defined above) were computed respectively.

To assess the statistical significance of the metric values for each neuron a random permutation test was used, using shuffled trial indexes (n = 1000). The p-value was computed as the fraction of shuffled indices that exceeded or equalled the observed value. Values were considered statistically significant when p < 0.05. If a neuron passed only the gOSI test, it was considered an orientation selective neuron. If a neuron passed the gDSI test or both tests, it was considered a direction selective neuron.

Optic flow stimuli and local drifting Gabor stimuli were used to confirm the response distributions observed with sinusoidal drifting gratings. For optic flow stimuli, direction selectivity was assessed similarly to drifting gratings. For local drifting Gabors stimuli, direction selectivity was assessed based on the presence of a receptive field. For each neuron, ETAs were computed separately for each Gabor directionality. The presence of a receptive field in a single ETA classified the neuron as direction selective.

#### Polarization degree selectivity metric

Medulla neuron event triggered average responses to polarized sparse noise (local DoLP decreases) were estimated as above, for both horizontally and vertical polarization angles. We then calculated the average spike rate of the neuron for 200 ms, triggered on times when a stimulus was present at the peak position of the event triggered average. Note that this does not require a receptive field to be estimated, as we wanted to include cases where a neuron was selective for single polarization angles. Spike rates were estimated by binning spike times across trials at 1 ms resolution, normalizing to Hz, and smoothing with a Gaussian filter (sigma = 5 ms). Spike rate vectors for horizontal and vertical polarization angles were then correlated (Pearson’s correlation coefficient) to determine the polarization degree selectivity metric, as done for the OL cortex.

#### Receptive field size of LFP

Local field potentials were high pass filtered at 0.1 Hz using a 4th-order Butterworth filter, then resampled at 1 kHz before bandpass filtering between 1 and 100 Hz with a 4th-order Butterworth filter. The event triggered average signal at each spatial position of the monitor during a sparse noise experiment (squares = 5×5 visual degrees, stimulus frequency 10 Hz) was then calculated for events of the square turning from gray to white, using a time period starting at event time and extending 125 ms. The mean over all channels of the event triggered average was subtracted to remove nonspecific signal, and the sum of the negative deflection (signal < 0) was taken as the LFP event triggered average response. LFP receptive fields were then estimated as in single unit event triggered average, described above but using a peak detection threshold of 4 SD of null, extending RF to where the ETA crosses 3 SD of null, and no Gaussian smoothing.

#### Contrast thresholds

To estimate the contrast thresholds of the medulla population, we combined neural recordings across multiple experiments responding to drifting gratings at a range of contrasts (light intensity or DoLP, see ‘Visual Stimuli’ above). Trials were binned by contrast (17 bins spanning −1 to 1 Weber contrast, central 15 used for DoLP stimuli, central ‘null’ bin centred at 0 contrast) and split into folds for cross validation (4 train and 1 test folds, leave-one-out). For each neuron, the spike rate function for each bin and fold was estimated by binning all peri-stimulus spike times at 1ms resolution, normalising to Hz, and smoothing with a Gaussian filter (SD = 5 ms).

To determine visually responsive neurons, spike rates were averaged across folds, and the last 200ms of each 1s trial (500ms stimulation, 500ms blank, see above) was subtracted as baseline. The response magnitude for each bin was taken as the mean absolute value of the difference from baseline firing, for the first 200 ms of each trial (onset response). Response magnitudes were smoothed with a Gaussian filter (SD = 1 bin), and neurons with a min-max difference in response magnitudes of over 2 Hz were considered responsive.

Responsive neuron spike rates by folds were used for SNR analysis. They were concatenated to form a population activity array of (folds x contrast bins x time x neuron), using the first 200 ms of each trial (onset response). Trials were averaged over folds for display (Fig. 5c). Response magnitudes were min-maxed normalised for single neuron reporting (Extended Data Fig. 14a-c).

For cross validated analysis of SNR, we divided data into train (4) and test (1) folds. Spike rates were normalized across neurons by z-scoring, using the mean and s.d. computed from train folds. We reduced the dataset dimensionality using PCA on the training set, retaining 70% of the training-folds variance and projecting all folds into PC space. For every contrast bin in the test fold, we calculated Euclidean distance between its activity vector and each null-contrast vector from the train folds. We then computed the signal to noise ratio (SNR) as:

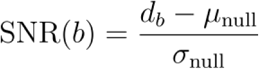

Where *d_b_* is the mean distance for bin *b*, *µ*_null_ is the mean across-fold distance of the null bin to itself, and *σ*_null_ is the across-fold s.d. of that distance. SNR was computed separately for every fold. Contrast thresholds were estimated by starting at the null bin and moving sequentially to higher (or lower) contrast bins. For each bin we applied a one-sided Wilcoxon signed-rank test across folds. The first bin whose mean SNR > 0 and p < 0.05 was taken as the neural population’s contrast threshold.

#### Intensity + Polarization stimuli analysis

combined intensity + polarization moving bar stimuli trials were first separated into 10 x 10 bins by contrast (intensity and DoLP). For each contrast bin, we estimated a neuron’s average spike rate by binning peri-stimulus spike times at 1ms resolution for a 1s time window following stimulus start time, normalizing to Hz, and smoothing with a Gaussian filter (SD = 5 ms). The last 100 ms of each contrast bin’s rate (baseline rate) was then subtracted to calculate stimulus response rate. A scalar response value for every contrast bin was calculated as the maximum of the absolute value of this response rate. This 10 x 10 response matrix was Gaussian smoothed (SD = 1 bin), and neurons with responses to any contrast bin above 15 Hz were min-max normalized before further population analysis (Fig. 5g,h).

For clustering, we used Leiden community detection (number of neighbours = 25, resolution parameter = 1) on a graph of neural responses. Responses across contrast bins were flattened into 100 dimensional vectors, and centred (z-scoring) before constructing a neighbourhood graph and running the Leiden algorithm. We utilized UMAP for visualisation of the results (n neighbours =15, min dist = 0, metric = ‘euclidean’).

To quantify how medulla neurons jointly encode light intensity and polarization degree information, we fitted an additive linear model to each neuron’s response matrix. Models were of the form:

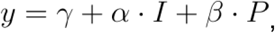

where *y* is the normalized response, *I* is intensity contrast, *P* is DoLP contrast, and *γ*, *α*, and *β* were fit by ordinary least-squares.

Because each response matrix was smoothed with a 1 bin Gaussian (σ = 1), adjacent bins are not independent. To avoid overestimation of the statistical significance of our model fit, we employed a tile bootstrap. 3×3 tiles of the 10×10 response matrix were resampled with replacement, and a linear model was fit to this shuffled data. This procedure was repeated 1000x, and 2.5 % and 97.5 % percentiles of the bootstrap distribution for each coefficient were taken as 95% confidence intervals. Statistical significance (p-values) were assessed as twice the fraction of sign changes between the bootstrap distribution for each coefficient and the original coefficient estimate. P-values were corrected for multiple comparisons using the Benjamini-Hochberg false discovery rate procedure (*α* = 0.05), with 99% (381/385) neurons showing significant coefficients (*α*, *β*, or both).

### Image processing for display

For display purposes, some of the light-sheet images were contrast enhanced using a gamma correction (γ = 2.0 in Fig. 3a; γ = 2.0 and 5.0 for nuclear staining and CM-DiI channel, respectively, in Fig. 3g, and Extended Data Fig. 9a-b; γ = 2.1 and 2.3 for nuclear staining and CM-DiI channel, respectively, in Extended Data Fig. 9c-f).

### Anatomical analysis

#### Medulla tree stereotypy

Light sheet volumes of nuclear stained brains were resampled at 10 µm isotropic voxels for analysis. Brains were warped to the atlas reference frame, and then a binary mask was applied to mark voxels belonging to the medulla. The medulla tree was then marked through image binarization. Two adaptive thresholds were used (Gaussian smoothing with 21 and 61 voxel sizes) to capture the range of spatial scales present, and the logical ‘or’ operation applied to make a binary mask. medulla tree similarity was calculated as the intersection over union (IoU) metric of all pairs of binarized brains A and B, within the medulla mask, as

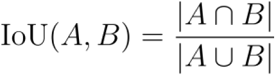

For display, Fig. 3c shows a slice of two registered brains sampled at 2.5 µm isotropic voxels.

#### Measuring neuropil-tree volume

Brains were registered and medulla tree binarized as above. Within the medulla, a distance map was computed measuring the Euclidian distance from the IGL. Binarized volumes were denoised using morphological opening of 2×2×2 voxels, and objects less than 10,000 voxels large were removed. The denoised binary volume was then skeletonized, and the local radius calculated using a distance transform from the skeleton to the binary volume. This was then converted into a local volume as V = 3/4πr^3^. For spatial averaging, The volume was averaged over non-overlapping 50×50×50 voxel cubes for plotting against distance. For curve fitting, a 5th order polynomial was fit to these distance-volume pairs.

#### DiI tracer analysis

Light sheet volumes of DiI injected brains were resampled at 10 µm isotropic voxels for analysis. The medulla was masked by inverse warping the brain atlas into the reference frame of each DiI injected brain. The injection electrode often left a trace of residual DiI, this was masked manually. DiI-labeled somata were extracted from these masked volumes through 2D difference of Gaussian filtering (SD = 1 µm and 50 µm) followed by global thresholding on all digital coronal slices of the volume. Binarized volumes were then computationally rotated to center the injection site and position the superficial brain (containing tracer spread) above this site. Injection sites and superficial brain points were selected manually. Volumes were translated to center the injection site. They were then upsampled 2x to avoid loss of data following rotation, and clipped to a cube 1000 µm/side to reduce computation time. Finally, volumes were rotated to align the vector connecting injection site and superficial brain point to the vector (0,1,0).

Tracer spread was visualised using a 2D polar histogram (angle bins=180, radial bins=36) of labeled voxels, using Euclidean distance in 3D as the radial distance from the central injection site, and X and Y angle as the 2D angular position. Histograms were normalized by dividing by the total values per radial bin, to show angular distribution insensitive to signal strength, which was strongest at positions closest to the injection site. Normalized histograms were then smoothed with a Gaussian filter (SD = 2 bins with ‘wrapping’ border).

### Neural process density

For constructing a histogram of neuron morphology (Fig. 3f), traced neurons were downsampled to 1 μm/pixel and rotated such that the superficial brain, inferred from the arrangement of cell islands in the brain slice, was set at 90°. A polar histogram was then constructed using 36 radial and 180 angular bins. The resulting histogram was smoothed with a Gaussian filter (std. 2 bins with ‘wrapping’ border).

### Field measurements

#### Underwater imaging surveys

All underwater imaging was conducted via SCUBA diving in compliance with the OIST Dive Safety Program and local regulations. Surveys were conducted at sites near the Okinawan coast; precise locations were selected based on logistical feasibility, including accessibility from shore, availability of depth profiles, consistent visibility conditions, and the presence of large, unobstructed, and relatively flat areas of seafloor. These included Koki Park North Beach (gps: 26.547248, 127.951244), Seragaki (gps: 26.506332, 127.881270), Apogama (gps: 26.498270, 127.841446), Seragaki-North (gps: 26.506373, 127.885829) and sites nearby. Depths ranged from 6-20 m, with tidal ranges of approximately 2 m.

We imaged in intensity and polarization a set of grayscale reference cards, positioned against a blue water background approximately 1-1.5m above the sea floor, at 10 m depth. Intensity images of the cards were collected along transects using a photographic single-lens-reflex camera (D5 Mark IV, Canon, Tokyo, Japan) equipped with a 100 mm macro lens, mounted in an underwater housing (Nauticam, Bournemouth, UK). All images were taken by keeping aperture, shutter speed and iso settings fixed across viewing distances spaced by 1 m. Polarization images of the greyscale standards at the same viewing distances were collected using a Division of Focal Plane (DoFP) polarization camera (Blackfly mono BFS-U3-51S5P-C, Teledyne FLIR, Wilsonville, USA) mounted in a custom-built underwater housing (SB Powell, University of Queensland, Australia), and using manually controlled aperture, exposure and gain. Stokes parameters and DoLP were calculated using standard formulae^82^.

A separate set of polarization images of white reference cards (30 × 22 cm) along transects in varying environmental conditions were collected using a digital photographic camera (TG-6, Olympus,Tokyo, Japan) equipped with a rotatable polarizing filter and stabilized by a tripod. At each location, a standardized set of nine images was taken at intervals of 3 seconds of the target, consisting of images at three polarizer angles (0°, 45°, and 90°) with three images per angle. ISO and aperture were fixed at 100 and f/2.0, respectively, and exposure settings were locked for the duration of each 9-shot sequence. Images from all three camera systems were captured exclusively during daytime under ambient natural light (no artificial illumination).

#### Transparent live prey imaging

Images of live transparent shrimp affixed to a stabilizing mount on a tripod were taken at 0.5 m distance, at 10m depth and identical locations to transect data. Locally caught wild shrimp specimens (multiple taxa), normally used to feed squid in the lab, were temporarily anesthetized prior to imaging using seawater cooled with ice to approximately 4 °C. This method induced immobility without releasing any chemicals into the marine environment or causing observable harm. Anesthetized individuals were gently clamped to the tripod mount and imaged within a 30–45-minute immobilization window. Long antennae were clipped, as they could not be stabilized underwater for imaging.

### Analysis of underwater images

For TG-6 camera data, sets of three images at each angle of polarization (horiz, diag and vert) were combined to make DoLP and intensity images. In the case of 9 shot sequences, the images were assembled as DoLP image 1: image 1,3,7, DoLP image 2: image 2,4,8, DoLP image 3: image 3,6,9. First, the green channel of the RAW Bayer array was extracted for analysis, with this channel approximating the spectral sensitivity of cephalopod rhodopsin^102^. For reference card imaging, small image shifts were removed by manually selecting the center of the white board in each image and translating all images to match the horizontal image. For shrimp images, images were aligned by detecting SIFT features^103^, brute force matching, computing the homography matrix from the matched features, and warping under a perspective transformation. Next, Stokes parameters were calculated from the three aligned images:

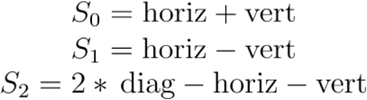

With DoLP and intensity images computed as:

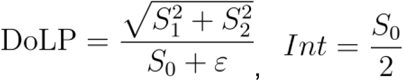

With *ε* = 1e-8 used for numerical stability. Visibility was quantified using Weber contrast calculated on intensity images, and ΔDoLP on polarization images. For every frame, a region of interest contained in the reference card/white square was taken as foreground (patch), and a region of the water column near the foreground was taken as background (bg). Foreground and background regions were averaged separately. Weber intensity contrast was computed as

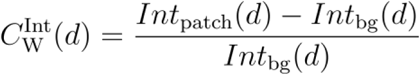

where *d* is horizontal distance. ΔDoLP was calculated as

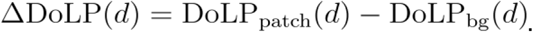

Sighting distances were estimated as the distance where an exponential fit to Weber intensity contrast or ΔDoLP crossed the detection threshold estimated from physiological data, up to the maximum distance measured (25m). To facilitate comparison between values, Extended Data Fig. 15c-l normalises (divides) by the respective detection thresholds.

### Use of Large-Language-Model assistance

We used ChatGPT (models ‘o1’ and ‘o3’, accessed from December 5, 2024), developed by OpenAI, to generate and format code for data visualization. All code, results and figures were manually reviewed and validated by the authors.

### Statistics and reproducibility

Unless stated otherwise, data are mean ± SD. For box plots: margins are 25th and 75th percentiles; middle line, median; whiskers, boundaries before outliers; outliers (+) are values beyond 1.5X interquartile range from the box margins. Experiments were repeated independently several times with similar results, with numbers of repetitions stated in text.

## Acknowledgements

We are grateful for the help and support provided by the OIST Cephalopod Research Support Team, the Scientific Computing and Data Analysis, Engineering, and Marine Science Station sections, Research Support Division at OIST, and the Harvard Medical School Electron Microscopy Facility. We thank Shoi Shi and Michael Hausser for comments on an earlier version of this manuscript, and all Reiter lab members for assistance and discussion. We thank Judit Pungor and Cris Niell for advice on cephalopod neurophysiology, and Tim Harris for assistance with Neuropixels probes. We thank Masatoshi Asato, Takenari Higa and members of the Nago Fishermen Union for providing wild squid eggs and boat space, Hiroki Touyama and the Onna Village Fishermen Union for providing wild squid breeders, Sam B Powell and N Justin Marshall for developing the DoFP polarization camera underwater housing and control system. This research was funded by the Okinawa Institute of Science and Technology, MEXT KAKENHI grants 23H02518 (SR), 24K18224 (TM), JP24H02309 (YG), JST grants JPMJFR224J and JPMJCR24T4 (SR). TM was funded by a Grant-in-Aid for JSPS Fellows (JSPS KAKENHI 21J01369). LM was funded by the Burroughs Wellcome Fund CASI award. LR was funded by the HHMI Hanna Gray Fellowship. MS was supported by the Marie Skłodowska-Curie postdoctoral fellowship 101066328 funded via the Engineering and Physical Sciences Research Council grant EP/X020819/1 and by the Young Researchers 2024 (MSCA) Grant, funded by NextGenerationEU.

## Author information

### Contributions

Project definition TM, KT, SR. Animal care: GDM, MHa, TLI. Imaging experiments: TM, YK. Electrophysiological experiments: KT, YK, MHi. Anatomical experiments and analysis: TM, KT, TTVD, RT, VG, LD, LR. Fieldwork: GDM, MS, MHo, MHi, KA. Data analysis: TM, KT, RT, DS, LM, SR. Project supervision: NB, MHo, YG, SR. SR wrote the manuscript with participation of all authors.

Corresponding author SR

## Ethics declarations

### Competing interests

The authors declare no competing interests

### Code availability

The code developed in this study will be posted in a repository on GitHub following publication.

### Data availability

Data are available from the corresponding authors on request. Sample datasets will be provided with the analysis code following publication.

## Notes

### Competing Interest Statement

The authors have declared no competing interest.

